# cellHarmony: Cell-level matching and holistic comparison of single-cell transcriptomes

**DOI:** 10.1101/412080

**Authors:** Erica AK DePasquale, Phillip Dexheimer, Daniel Schnell, Kyle Ferchen, Stuart Hay, Íñigo Valiente-Alandí, Burns C. Blaxall, H. Leighton Grimes, Nathan Salomonis

## Abstract

To understand the molecular pathogenesis of human disease, precision analyses to define molecular alterations within (and between) disease-associated cell populations are desperately needed. Single-cell genomics represents an ideal platform to enable the identification and comparison of normal and diseased transcriptional cell states. We note that disease-associated perturbations usually retain cellular-identity programs (core genes), providing an appropriate reference for secondary comparison analyses. Thus, we created cellHarmony, an integrated solution for the unsupervised analysis and classification of cell types from diverse scRNA-Seq datasets. cellHarmony is an automated and easy-to-use tool that efficiently matches single-cell transcriptomes using a community clustering and alignment strategy. Utilizing core genes and community clustering to reveal disease and cell-state systems-level insights overcomes bias toward donor and disease effects that can be imposed by joint-alignment approaches. Moreover, cellHarmony directly compares cell frequencies and gene expression in a cell-type-specific manner, then produces a holistic representation of these differences across potentially dozens of cell populations and impacted regulatory networks. Using this approach, we identify gene regulatory programs that are selectively impacted in distinct hematopoietic and heart cell populations that suggest novel disease mechanisms and drug targets. Thus, this approach holds tremendous promise in revealing the molecular and cellular origins of complex diseases.

## INTRODUCTION

Single-cell RNA-sequencing (scRNA-Seq) provides the unique ability to profile transcripts from diverse cell populations along a continuum of related or disparate states (Olsson et al. 2016). In addition to defining known and novel cell populations, single-cell technologies can identify disease-related gene regulatory programs which underlie molecular and cellular dysfunction. While diverse single-cell experimental platforms exist to facilitate such analyses, there is an urgent need for integrated and easy-to-use computational approaches to identify discrete differences between comparable diseased and healthy cells. Given that most tissue scRNA-Seq analyses will potentially identify dozens of cell populations, such an exercise becomes non-trivial, as distinct cell populations will have different transcriptional, cellular, pathway, and gene regulatory network impacts. Furthermore, cellular and molecular differences can occur in either a cell-state-specific manner or across a spectrum of related cell states, requiring new holistic analysis solutions. Given the complexity of the analyses required to achieve these goals, automated solutions that can be applied by both experienced bioinformaticians and conventional laboratory biologists are ultimately required.

The development of workflows to provide disease-level insights requires reproducible mapping and comparison of single-cell transcriptomes across one or more samples. Recently, several approaches have been developed to align transcriptionally similar cells within the same platform, across distinct platforms, and even across species. These methods include approaches designed to propagate cell and cell-type labels across datasets, such as scmap, and joint alignment methods that simultaneously align similar cells into to common or distinct clusters independent of batch, donor, or other technical effects, such as Biscuit, MNN, and Seurat CCA (Prabhakaran et al. 2016; Butler and Satija 2017; Haghverdi et al. 2018; Kiselev et al. 2018). Existing label projection methods begin with identification of variable genes in a reference scRNA-Seq dataset and align query cell profiles to these references, using a basic or approximate nearest neighbor search. While the joint analysis of scRNA-Seq datasets can yield cell-population insights and cell-type relationships (e.g., naïve versus regulatory T-cells), the alignment of distinct scRNA-Seq datasets into a shared subspace can result in unintended artifacts. In particular, significantly-perturbed gene expression arising from disease, drug treatment, or environment effects can result in imperfect alignments that result in dataset-specific skewing of cell populations. The end result of joint-alignment approaches can be the formation of clusters that are apparently unrelated to normal cell states. Thus, failure to produce clusters of equivalent cells might reveal a bias toward donor and disease effects imposed by joint-alignment. While tools such as Seurat enable the direct comparison of cells within the same population across conditions (differential expression analysis), iterating this analysis over potentially dozens of cell populations, integrating these results, and obtaining systems-level insights remains an unsolved challenge.

Herein, we describe a new approach called cellHarmony, which provides the unique ability to obtain a unified systems-level view of molecular, cellular, pathway, and network-level differences between all aligned query and reference cell populations in an automated manner. We observed that in comparable disease and normal datasets, the cellular-identity programs (core genes) are retained; providing an appropriate reference for secondary comparison analyses. To improve the accuracy of cell projections, cellHarmony creates a k-nearest neighbor graph of each sample to match similar sample communities (Louvain clustering) and then directly match the cells within those communities. This strategy enables the fast alignment of similar small communities without biasing the alignment to outliers. When compared to joint-alignment methods and other label projection approaches, cellHarmony results in improved alignment accuracy and is less sensitive to genetic, experimental and technological biases. Once aligned, clusters or cell-type names from one dataset can be mapped onto another and placed within a continuum of differentiation. These data are integrated and jointly visualized in multiple formats. To identify the precise impact of genetic, chemical, disease, or other perturbation, cellHarmony extracts cell-state-specific differences (cell frequency and gene expression), finds, organizes, and visualizes co-regulated cell populations, examines the pathway-level impact on distinct gene modules, and produces putative gene-regulatory networks based on prior knowledge. Through seamless integration within AltAnalyze (Emig et al. 2010; Olsson et al. 2016), users can jointly perform the unsupervised analysis of extremely large scRNA-Seq datasets using the algorithm Iterative Clustering and Guide-gene Selection (ICGS, version 2) prior to alignment, with little to no required bioinformatics expertise (Venkatasubramanian et al. 2019). Automation becomes a necessary step in these analyses, where dozens of cell populations are likely present in large single-cell RNA-seq datasets. Using this approach, we are able to effectively derive important novel cellular, disease, and gene regulatory insights when applied to diverse cell atlases (mouse and human) and disease datasets (leukemia and heart failure).

## METHODS

### Algorithm Description

#### Implementation and Requirements

cellHarmony is compatible with Python 2.7 and is distributed as a component of the software AltAnalyze (https://github.com/nsalomonis/altanalyze) and separately as dedicated python source code for the community clustering alignment algorithm (https://github.com/AltAnalyze/cellHarmony-Align). AltAnalyze is an easy-to-use data analysis toolkit that provides both automated workflows (e.g., RNA-Seq raw data processing, alternative splicing, scRNA-Seq unsupervised analysis, microarray analysis) and separate à la cart analyses through the command-line and a dedicated graphical-user-interface. This code is supported for Linux, Mac, Windows, using computers with a minimum of 8GB of RAM (16GB+ recommended). Additional documentation, optional R pre-processing scripts (for Seurat input references), and tutorial videos are available from: http://www.altanalyze.org/cellHarmony. When run through AltAnalyze, cellHarmony works seamlessly with the embedded unsupervised analysis software, Iterative Clustering and Guide-gene Selection (ICGS) version 2 as input (Olsson et al. 2016; Venkatasubramanian et al. 2019). cellHarmony can be run both on the command-line and through the AltAnalyze version 2.1.1 graphical user-interface (Additional Analyses Menu > Cell Classification Menu). Pre-compiled graphical user-interface distributions are provided from http://altanalyze.org and the command-line version from GitHub or via installation from PyPI. Details regarding required and optional input files and additional information regarding the use of cellHarmony on a computational cluster, are provided in **Supplemental Information**.

#### Cell-Alignment from Community Clustering

To identify equivalent cell-types or cell-states from two independent scRNA-Seq datasets, cellHarmony employs a community clustering strategy to produce a network graph and define communities in both the reference and query dataset. In short, this approach initially matches communities of cells across samples and then selects the closest matching reference cell for each query cell, where a single reference cell can match to multiple query cells. This function performs the following steps:

##### 1) Define communities (partitions) within each of the datasets

Prior to defining communities, each dataset is restricted to the cell-population specific marker genes previously defined for the reference. When using ICGS version 2 unsupervised results (default), these markers are the top 50 cluster-specific marker genes for each cluster. The query and reference files are imported as Cell Ranger (10x Genomics, version 1.0-3.0 supported) sparse matrices (h5 or mtx) or as tabular input files (counts or scaled). By default, cellHarmony uses the ICGS-NMF produced output heatmap tab-delimited text file as its reference (**Supplemental Information** - Input Data). Utilization of each of these inputs produces equivalent downstream results. From this sparse matrix data, the program identifies the k-nearest neighbors (k =10 by default) for each cell using the python package Annoy (Aumüller et al. 2018) and creates a graph of these neighborhoods using the networkx python package. Once created, Louvain clustering is performed with the lowest possible resolution to find maximal partitions (r = 0). We denote the resulting communities of reference sample cells by {Cr_1_, Cr_2_, …, Cr_s_} and {Cq_1_, Cq_2_, …,Cq_t_} for the query sample communities.

##### 2) Find the closest matching communities between the query and reference

For each community in the reference sample (Cr_i_), a centroid mCr_i_ is calculated from the constituent cells using a simple mean for each gene and likewise mCq_i_ for each query sample community (mCq_i_). All pairwise similarities r_ij_ (numpy corrcoef function) between query (mCq_i_) and reference (mCr_i_) community centroids are computed and the most highly correlated (Pearson coefficient) reference community for each query community is identified. We represent result of this matching process for a query community as {Cq_j_:Cr_i_; ∀j, i:max(r_ij_)}.

##### 3) Find the closest matching reference cell for each query cell and propagate the label

The gene expression of each cell within the query community of a pairing {Cq_j_:Cr_i_} is compared (Pearson correlation coefficient) to each cell in the matched reference community Cri to find the closest match. The original cluster label (i.e., user-supplied label or ICGS/Seurat cluster number) assigned to the closest match reference cell is then propagated to the query cell. A new heatmap is created wherein every query cell is placed adjacent to its closest match reference cell; reference cells being grouped by the original clustering (see Input Data). However, query cells with a Pearson alignment correlation coefficient lower than the user defined threshold (default = 0.4) are excluded from downstream combined data visualizations, differential expression, and system level prediction analyses. The result is a set of aligned cell populations, each cell population consisting of all reference cells with a common label and any query cells with that label. We denote the set of aligned cell populations by {A_k_, k=1…K} where K is the number of ‘original’ clusters in the reference dataset. The relative frequency distribution of the (aligned) populations within the reference and query samples is compared using a series of K Fisher’s exact tests. Alternative options for running these functions and performance are described in the **Supplemental Information**.

#### Differential Expression Analysis

To identify impacted genes, networks, and cellular processes across cell-populations, cellHarmony performs a differential expression (DE) analysis between each aligned query and reference cell population {A_k_, k=1…K}. Differential expression is performed using an empirical Bayes moderated t-test with Benjamini-Hochberg (BH) adjustment separately for each comparison. An overall moderated t-test for each gene comparing the pooled query cells to pooled reference cells is also performed, with the BH correction applied. We denote the adjusted p-values from these analyses by {p_1_, p_2_, …p_K_} and p_O_ for the overall test. The default threshold for differential expression is fold > 1.5 and p < 0.05 (FDR corrected). Users can modify these thresholds within the graphical user interface or from the command-line (**Supplemental Information**). This process is automated for cell populations in the two compared datasets that share the same provided or assigned labels between the reference and query, respectively. The K pairwise query-to-reference comparisons are performed for cell populations in which at least 20 cells are present from both the query and reference datasets. For datasets with fewer than 200 total cells (i.e. Fluidigm C1), this requirement is relaxed to 4 cells in each aligned cluster. The results are saved to tab-delimited text files with summary statistics and basic annotations, along with summary graphical outputs, and are used for downstream systems-level analyses.

#### Systems-Level Predictions

In comparisons where a molecular, genetic, or chemical perturbation results in abnormal cell biology, cellHarmony can be used to identify which cell populations are principally impacted in both cell frequency and gene expression. By examining the differences that are shared among different cell states and those that are unique to a specific state, cellHarmony enables the determination of holistic pathway and gene regulatory network impacts. Specifically, cellHarmony determines whether gene expression differences across cell-states are global (impacted in most cell populations), local (i.e., cell-state specific), or co-regulated/regional (i.e., specific to a subset of cell clusters). Note that although some clusters can be excluded from the differential expression analysis due to insufficient cell population size (see above), we continue using the notation for the full set of aligned clusters {A_k_: k=1…K}.

To assess global gene up- and down-regulation, the differential expression analysis is repeated for all query cells compared to all reference cells, regardless of the cluster to which they are aligned. Only genes for which p_O_ < 0.05 and p_i_ < 0.05 for at least 2 {i:1…K}, with consistent direction of fold change, are considered as candidates (see below) for the globally upregulated or globally downregulated group, depending on the direction of the fold change. Genes that meet the DE criteria stated above *for only one cell population* form the local DE groups (i.e. p_i_ < 0.05 for only one {i:1…K}) represent cell-state specific profiles. To define additional predominant patterns of co-regulation (common effect among multiple but not all clusters) for each gene, the cell-state specific p-values comparing each query to reference cluster (A_k_) are collected into a vector of length K. The vector is recoded to entries of 1, −1 or 0 as follows: 1 if p < 0.10 & sgn(logfc) = 1, −1 if p < 0.10 & sgn(logfc) = −1, 0 otherwise to indicate upregulation (1), downregulation (−1), or no significant alteration (0). The p-value threshold of 0.10 was chosen to provide increased sensitivity in clusters with small numbers of cells. For example, a gene with pattern [0,1,0,1] would represent co-regulated DE for clusters 2 and 4. The four most frequently occurring patterns of co-regulation (excluding global and cell-state specific patterns) are selected for an additional round of differential gene expression analyses, comparing query to reference cells in the resulting aggregated cells clusters. For example, if the top pattern is [1, 1,0,0], a test of pooled clusters 1 & 2 in the query will be compared to cluster 1 & 2 in the reference scRNA-Seq dataset, yielding p_(co-reg)_. Each gene from the cell-state specific comparisons (local), is then assigned to the specific comparison group (global, local, or co-regulated) in which they are most altered, based on the smallest p-value of all comparisons, i.e. among {p_i_: i=1…k, p_O_, p_(co-reg)_} (additional criteria for global regulation noted above). The final genes are limited to significant genes in the first cell-state differential analysis, excluding new genes identified in the global or regional analysis, to prevent bias due to cell-type frequency variation in the query and reference (e.g., more B cells in query versus reference, highlighting B-cell upregulated gene expression).

To produce a unified representation of these differences, which can consist of dozens of differential expression comparisons, cellHarmony creates a combined heatmap with genes assigned to specific comparison groups. The cells in the heatmap are ordered by the original cell populations, with the genes ordered according to global, co-regulated, and local categorizations (p-value ranked), with upregulated and downregulated genes displayed as adjacent clusters. A gene set enrichment analysis is performed on the resultant gene clusters with the Pathway Commons database (human and mouse) or Gene Ontology (other species) (Ashburner et al. 2000; Cerami et al. 2011). Putative gene regulatory relationships in these gene modules are predicted using a second gene set enrichment analysis with the TRRUST, PAZAR and Amadeus databases (combined) to identify likely upstream transcriptional regulators and highlight clustered embedded transcription factors (Zambon et al. 2012; Han et al. 2018). Finally, these gene regulatory networks are further visualized using the ‘igraph’ pathway library and these same transcription factor target databases for each comparison.

### Statistical Methods for Comparative Analyses

To evaluate the agreement in label assignments for different label projection methods and for the software Reference Component Analysis (RCA) (Li et al. 2017), we used the Adjusted Rand Index (ARI) as proposed by Hubert and Arabie (Hubert and Arabie 1985) using the R package ‘clues’ (Wang et al. 2007). Confidence intervals (95%) for ARI were computed using the normal approximation as implemented in the R package ‘CrossClustering’ (Wang et al. 2007). Accuracy is assessed as the proportion of correct assigned labels for a given alignment algorithm, where perfect accuracy is defined as matching aligned and author assigned labels.

### Evaluation Datasets

See the **Supplemental Information** for details regarding the software inputs and outputs, heart scRNA-Seq produced in these studies, external validation datasets used, external software evaluation, data processing and analysis parameters, along with user guidelines for cellHarmony parameter tuning.

## RESULTS

The cellHarmony pipeline was created to rapidly and accurately align comparable scRNA-Seq datasets to identify the global, regional and local molecular impacts of diverse perturbations. For this purpose, cellHarmony uses a graph-based strategy to produce disjoint networks from a clustered single or combined reference against an un-clustered query (**Fig. 1A,B**). Alignment through community clustering provides three important advantages in the alignment of different scRNA-Seq datasets. First, individual cells can be quickly matched between large scRNA-Seq datasets by finding the most similar communities between datasets prior to matching individual cells in those partitions. Second, the produced alignments will be more stable than individual cell alignments which are inherently biased towards outlier cells. Finally, the alignment to cell centroids is highly reliant on the final cluster definitions, in which sub-clusters may be present or clusters inappropriately aggregated. Cell populations and cluster labels in the reference can be automatically assigned using the embedded unsupervised algorithm Iterative Clustering and Guide-gene Selection (ICGS) (Venkatasubramanian et al. 2019), which has been consistently shown to identify transitional cell-states (default ICGS Guide3 output) and exceedingly rare cell states from large scRNA-Seq datasets (ICGS-NMF output) (Olsson et al. 2016; Churko et al. 2017; Magella et al. 2017; Yáñez et al. 2017; Hay et al. 2018; Lu et al. 2018; Venkatasubramanian et al. 2019). While reference cell alignment is an important independent objective in many studies (i.e., developmental ordering of cells, cell-type identification), cellHarmony goes further by producing systems-level models of cellular and transcriptional differences in discrete and co-regulated cell populations. To achieve this aim, the software implements a unique comparison and aggregation approach to identify impacted gene and regulatory programs that may be restricted to individual cell-states, shared more broadly between specific states, or that are common to all populations (**Fig. 1C-F**).

**Figure 1.**
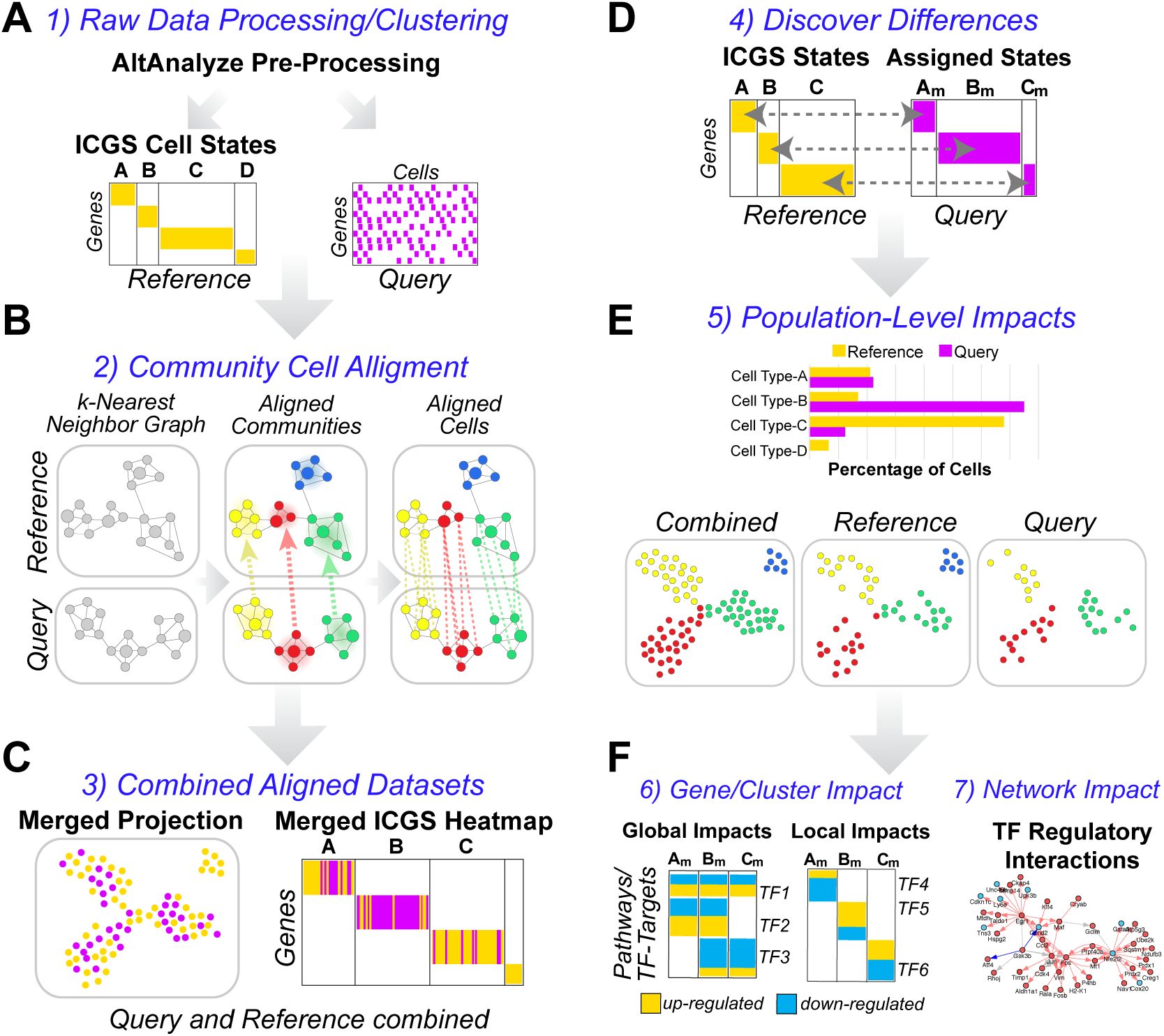
Integrated and automated holistic discovery of cell-state specific perturbations. Illustrated automated workflow of the cellHarmony analysis pipeline. A) Small and large (>100k cell) scRNA-Seq datasets can be pre-processed in the parent software AltAnalyze (expression scaling, outlier exclusion) for the query and reference dataset. Raw data can be input from multiple file formats (Methods). Both query and reference can be individual samples or combined collections (joined and analyzed or merged post unsupervised analysis with the cellHarmonyMerge function). The default unsupervised clustering population prediction method in AltAnalyze is ICGS version 2.0 (ICGS-NMF output), which provides different input options (population marker genes or guide-gene correlated results). B) Alignment of individual cells from the query to the reference is accomplished through community clustering of each dataset, following the creation of a k-nearest neighbor graph in each dataset. Prior to matching cells, Louvain partitions (cluster centroids) are matched between datasets to define the nearest neighborhoods (Pearson coefficient). Labels provided for the reference clusters (stem cell, macrophage…) can be provided as alternatives to cluster labels (c1,c2,c3…), based on the ICGS cluster cell-type predictions (curated by the user). C) Query cells that meet a minimum correlation threshold (Pearson coefficient > 0.4, default) are placed adjacent to their aligned reference cell in the ICGS heatmap and projected into a common UMAP coordinate space to determine the mixing of cells and qualitative transcriptomic differences. D) Quantitative transcriptome differences between query and reference cells are computed using a multi-step differential expression analysis between matched cell-states and E) population proportion differences reported (Fishers Exact test). F) Gene-level statistical differences are used to identify the most similar impacted cell-states (co-regulated), defined by comparing the frequency of genes similar patterns of regulation from a binarized p-value and fold-change matrix. Genes with expression best described by global regulation (impacted across all cell-states), co-regulated clusters (regional) or that are most restricted in their differences to specific cell states (local) are ordered statistically in a combined heatmap (left). Integrated with this heatmap are statistically enriched pathways and transcription-factor targets to determine the perturbation-specific impacts along a continuum of related and distinct cell populations. Putative transcriptional regulatory and curated protein-protein interaction networks are derived from each cell-state or co-regulated comparison to identify likely regulators and their targets among these impacted populations (right).

### Cell-alignment performance

The two principal approaches for comparing scRNA-Seq populations are: 1) those that propagate labels from one scRNA-Seq dataset onto another and 2) those that perform joint-alignment of cells from different datasets. An important conceptual advantage of the latter is that in principle datasets can be combined regardless of donor, batch, or disease effects through their joint analysis. However, when sample differences are sufficiently large, significant biases in the cell population predictions can result, as shown in the joint analysis of two distinct scRNA-Seq bone-marrow donor samples from the Human Cell Atlas (HCA) project (Hay et al. 2018) (**Fig. 2A** and **Fig. S1A,B**). This underlying problem can make the direct comparison of reasonably equivalent populations in different samples problematic. Label projection largely negates this issue, by finding the most similar equivalent cells in a query sample relative to the reference. This is demonstrated with application of cellHarmony to the same HCA data as illustrated by improved overlapping cell-type predictions and increased proximity within the UMAP projection (**Fig. 2B** and **Fig. S1C-E**).

**Figure 2.**
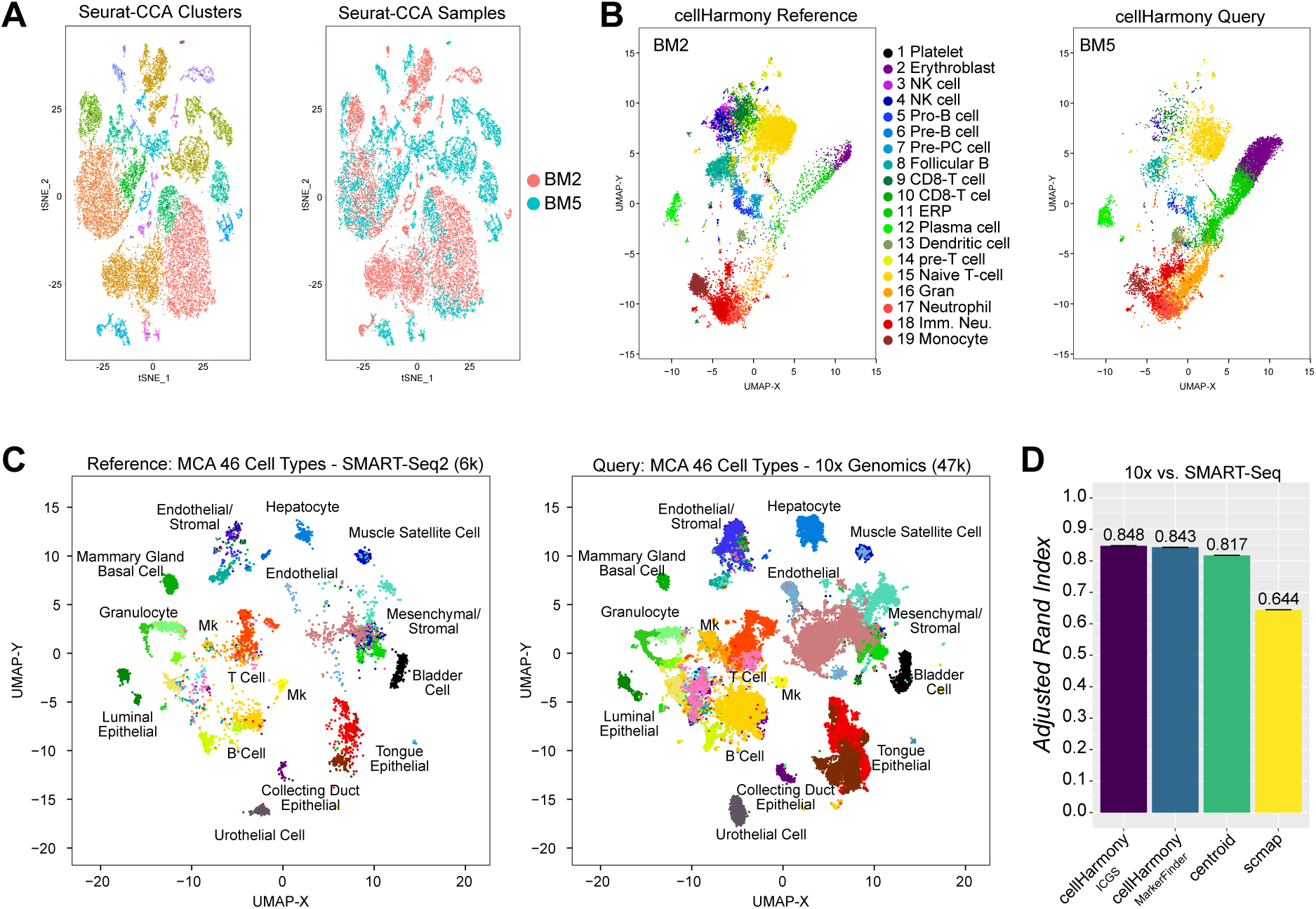
cellHarmony produces accurate alignments across technologies and diverse samples. A) An example of how transcriptomic differences can confound the identification of matching cell populations from scRNA-Seq. In this example, joint-alignment between two bone marrow donor scRNA-Seq datasets from the Human Cell Atlas project was performed using the Seurat CCA algorithm (version 2.3.4) and the data viewed as a t-SNE plot by cluster (left) or donor (right). B) UMAP projection of the same data from panel A following label projection in cellHarmony. The left plot shows cells only from the reference (BM2) and the right only from the query (BM5), colored by cellHarmony assigned clusters. Original cluster labels were assigned in AltAnalyze (GO-Elite gene set enrichment) for the ICGS-NMF produced gene clusters, using the gene-sets provided by Hay et al. C) UMAP projection of cells from 12 Tabula Muris tissues profiled using two different single-cell methods, SMART-Seq2 (reference, left) and 10x Genomics (query, right). Cells were restricted in both sets to those with common assigned Cell Ontology annotations by the study authors to allow for evaluation of the alignment agreements. D) Comparison of the cell-label assignment by three independent algorithms (cellHarmony, scmap, centroid-correlation) using the cells and labels from panel C. Results from cellHarmony are reported using variable genes derived from a supervised analysis of the Cell Ontology labels (MF=MarkerFinder) or from an ICGS analysis of all SMARTSeq2 cells. Adjusted Rand Index is reported as the overall measure of agreement to the author provided Cell Ontology labels with confidence intervals for ARI computed using the normal approximation.

To specifically evaluate the performance of this method relative to alternative label projection methods, we applied cellHarmony, scmap, and a centroid-based k-nearest neighbor alignment approach to a large mouse cell atlas dataset (Tabula Muris) with previously defined Cell Ontology classifications (n=46). This dataset includes 12 matched tissues profiled using two complementary scRNA-Seq technologies (10x Genomics and SMART-Seq2) and capture methods (unbiased versus Flow Cytometry). Although the two scRNA-Seq datasets generated by the Tabula Muris consortium were produced using independent single-cell methodologies, UMAP and heatmap projections of cellHarmony-aligned datasets indicate highly overlapping cell-state projections for the major cell populations, indicating that these core gene cell-type programs are highly reproducible (**Fig. 2C** and **Fig. S2A,B**). Using the Adjusted Rand Index (ARI) for alignment of the 10x Genomics query to the SMART-Seq2 reference to quantify similarity, community-based alignment in cellHarmony had improved similarity to the author-defined ground-state truth (ARI=0.85) with relatively high accuracy (83%), relative to centroid-based alignment (ARI=0.82, accuracy=80%) and scmap (ARI=0.64, accuracy=71%), without threshold-based filtering (**Fig 2D**). While an additional method, Reference Component Analysis (RCA) (Li et al. 2017), produced a similar ARI to cellHarmony (ARI=0.85), this centroid-based classification method does not perform label projection and cannot be used to assess accuracy (**Fig. S2C**). Effectively no differences were observed for cellHarmony when using markers obtained from two independent methods (MarkerFinder algorithm in AltAnalyze or ICGS). Filtering of query cells with the lowest percentiles (1%, 2%, …, 10%) of cellHarmony assigned correlations, indicates that cellHarmony predictions are relatively stable with more aggressive filtering (**Fig. S2D**). We find that mis-classifications from cellHarmony could largely be explained by alignment of cells with same or highly related cell-types across tissues (e.g., Muscle Macrophage as Trachea blood cell) (**Fig. S2E**). To determine the thresholds at which cellHarmony predictions fail to be accurate, we excluded two tissue-restricted cell populations that clearly separated out from other cell-types in the UMAP graph, liver hepatocyte and kidney collecting duct, and classified those cells back into the filtered reference dataset. These data indicate that cellHarmony is more likely to produce true positive alignments above a Pearson correlation >0.3, with alignments below that range indicating likely false positive associations for cells from the same technological platform (**Fig. S2F**).

In the mouse cell atlas dataset, it is important to note that there are far more discrete cell populations expected than the annotated 46 cell types. Notably, application of the cellHarmony Merge function to the 12 tissues profiled with the 10x Genomics platform, identified 171 transcriptionally distinct cell populations (rather than the reported 46), where similar cell-states were merged within and between tissues (Methods) (**Fig. S3A**). Dividing the 10x Genomics dataset in to a random training and test set again ranked cellHarmony highest for ARI, followed closely by scmap and centroid-based classification (**Fig. S3B**). Hence, cellHarmony is accurate across technological platforms to yield high precision cell-state alignments.

### Cell-population and gene-level impacts in aligned transitional cell states

To obtain higher order insights, cellHarmony requires an effective method to determine gene expression differences when comparing query cells to reference cell-states. For such comparisons, it is expected that the end-user identifies any possible batch-effects and corrects for them as needed, prior to performing cellHarmony. As an initial disease evaluation, we compared a previously described Fluidigm scRNA-Seq dataset of 382 murine bone marrow hematopoietic progenitors (BM) (Olsson et al. 2016) to genetically perturbed hematopoietic progenitors derived from a mouse-model of human Acute Myeloid Leukemia (AML) carrying both Flt3-ITD and Dnmt3a mutations (Meyer et al. 2016). Splenic c-kit positive AML cells from these animals were aligned to the c-kit positive bone-marrow-progenitor reference from the same strain of animals, collected from the same laboratory using the same scRNA-Seq methodology (Fluidigm C1). These reference cell populations include experimentally validated transitional states, notably: 1) a relatively frequent group of cells with myeloid, erythroid and megakaryocyte coincident expression (Multi-Lin), 2) progenitors in a metastable uncommitted state (GG1) and 3) bi-potential intermediates that produce specified monocytic or granulocytic progenitors (IG2). The transcriptomes of the aligned AML cell-populations were qualitatively matched to those of the wild-type cells, suggesting the core cellular gene program of these cell-types are retained (**Fig. 3A**). These AML cells were aligned with the highest frequency to wild-type dendritic cells, monocyte progenitors and IG2 cells (**Fig. 3B**). Hence, these data suggest that a genetically defined subset of AMLs may derive from a short-lived cellular intermediate with bipotential monocytic and granulocytic potential (IG2).

**Figure 3:**
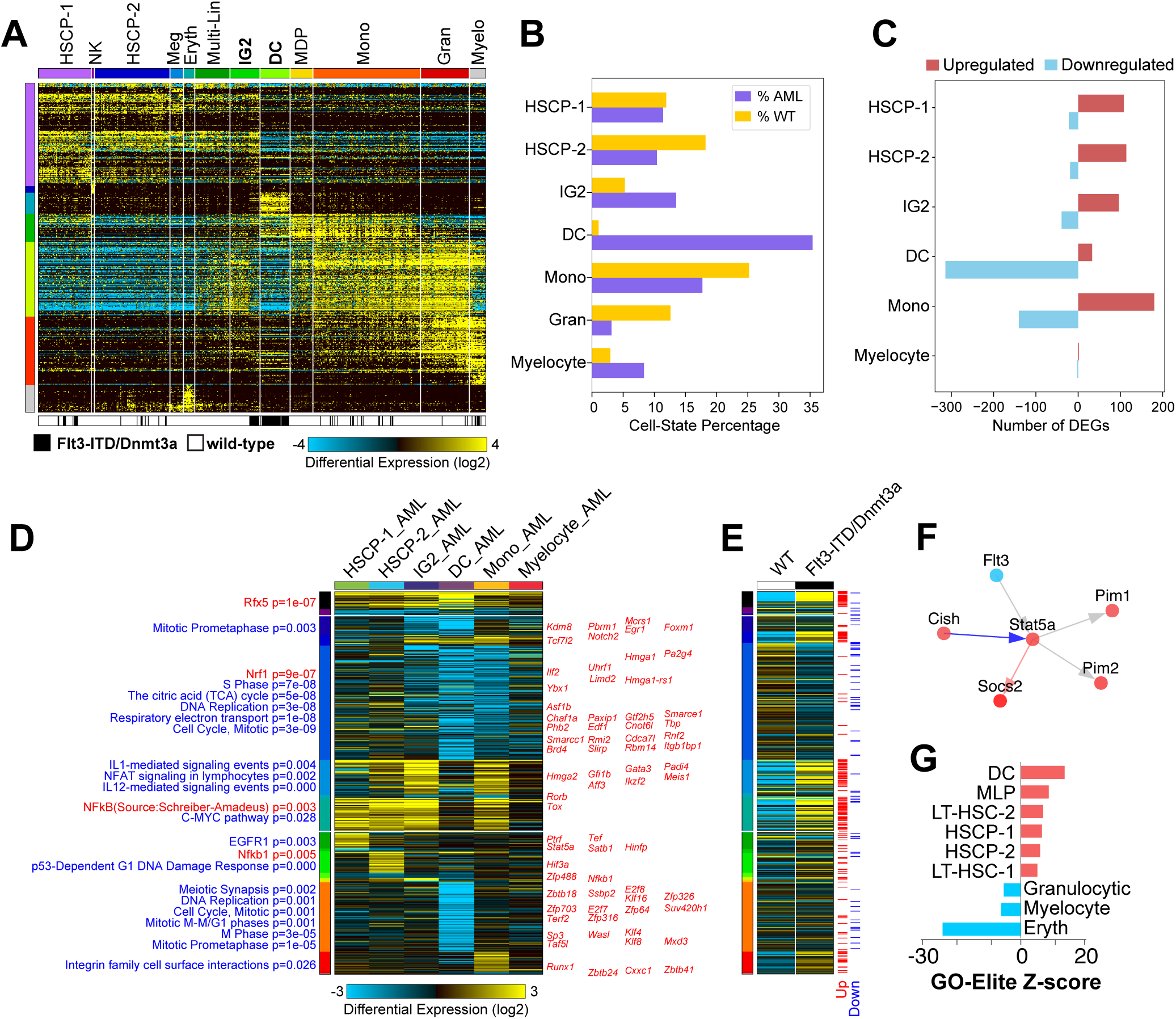
Cell-state specific impacts in transitioning progenitors and AML. A) cellHarmony alignment of previously profiled splenic c-kit-positive AML cells relative to ICGS results from all captured normal mouse bone marrow hematopoietic progenitors (reference sample, Olsson et al. 2016). B) AML cells were aligned most frequently to annotated dendritic cells (DC), monocyte progenitors (Mono), and previously described bipotential hematopoietic progenitors (IG2, Olsson et al. 2016). The AML DCs were high enriched relative to wild-type DCs by cellHarmony Fisher Exact test (p=4.8E-24). C) Number of differentially expressed genes (DEG) by cell-state (upregulated and down-regulated) for AML versus wild-type cell populations (fold>2, p<0.05, FDR adjusted). Negative numbers on the x-axis indicate downregulation (i.e. “-100” means “100 downregulated genes”). D) Heatmap of AML vs. wild-type fold changes for all significant differentially regulated genes from panel C. Genes are grouped as global, regional, and local transcriptomic differences (DEG p-value ranked), with DEGs demonstrating global regulation shown at the top of the heatmap, co-regulated cluster impacts in the middle, and cell-state specific impacts at the bottom. The regional or co-regulated clusters indicate shared patterns of gene expression across multiple cell-states (e.g., HSCP-1 + HSCP-2 + IG2 + Mono). For each pattern, a separate up-regulated and down-regulated cluster is shown. Known transcription factors are displayed to the right of the heatmap where present by cellHarmony and enriched Pathway Commons gene-sets (blue) and transcription factor target sets (red) are displayed on the left next to each cluster in ranked order of significance (bottom to top). E) Corresponding gene expression differences from bulk RNA-Seq of the AML model compared to wild-type bone marrow controls as an independent verification. Red lines = upregulated genes (fold>1.5, empirical Bayes moderated t-test p<0.05 (FDR adjusted)), Blue lines = downregulated genes (fold<-1.5, empirical Bayes moderated t-test p<0.05 (FDR adjusted)). F) Core predicted cellHarmony gene regulatory network for HSCP-2 AML versus HSCP-2 wild-type. Red nodes = upregulation, blue nodes = downregulation, red arrow = transcriptional regulatory interaction, blue arrow = inhibitory interaction. G) Validation of single-cell proportional differences. Statistical enrichment of cell-specific gene-sets from the software GO-Elite, of bulk AML RNA-Seq versus wild-type for cell-type specific gene-sets (BioMarker database).

For differential expression analysis, cellHarmony applies an empirical Bayes (eBayes) moderated t-test method. Compared to alternative algorithms, we find this approach to be effective at identifying cell-type-specific differences in gene expression in scRNA-Seq relative to deeply sequenced bulk RNA-Seq samples (**Fig. S4A**). Differentially expressed genes calculated from the comparison of the AML cells to their cellHarmony-matched populations indicate significantly differentially expressed genes in each cell population, with the majority of regulated gene programs shared between distinct combinations of cell-states, rather than globally perturbed across all cell-states (**Fig. 3C,D**). In particular, this analysis finds down-regulation of S-phase cell-cycle and TCA-cycle genes in dendritic and monocytic progenitors, upregulation of interleukin and NFAT signaling in IG2 and monocytic progenitors, but coordinated upregulation of a C-MYC regulatory program in the earliest progenitors (HSCP-1, HSCP-2, IG2) and monocytic cells. Visualization of differentially expressed transcription factors (TFs) within this representation further highlights the mis-expression of TFs in this model of AML. For example, the HSCP-marker genes *Gfi1b, Meis1, Gata3, Ikzf2* were upregulated in AML IG2 and monocytic populations, where they would normally be downregulated during differentiation (Olsson et al. 2016). These genes were previously shown to be aberrantly expressed in human AML (Gao et al. 2015; Liu et al. 2017; Thivakaran et al. 2018; Park et al. 2019). To verify these findings, we examined the same genes from the cellHarmony-ordered differentially-expressed-gene heatmap in bulk RNA-Seq from c-Kit+ AML splenocytes and wild-type c-Kit+ bone-marrow progenitors (**Fig. 3E**). This analysis confirmed much of the global, regional (HSCP-1 + HSCP-2 + IG2 + monocytic) and local expression differences (monocytic, dendritic, HSCP) when restricted to these genes. Next, cellHarmony analysis highlighted putative gene and regulatory interactions, including recently-identified *FLT3-ITD-*induced genes which can serve as drug targets (*Pim1, Pim2*) (**Fig. 3F**) (Green et al. 2015). As bulk RNA-Seq can reflect changes in cell frequency and/or cell-state-specific gene-expression differences, we analyzed all differentially expressed genes from the bulk analysis (**Fig 3G**). As expected, these data show an enrichment in gene sets for dendritic cells among upregulated genes and erythroid progenitors in down-regulated genes, consistent with the increased and decreased frequency of these aligned AML cell types, respectively. This suggests that the captured cell types were representative rather than randomly isolated.

### Computing disease-level impacts in diverse tissue cell-types

Our analysis of a murine AML model highlights the ability of cellHarmony to identify cell-state specific changes that are uniformly reflected from bulk RNA-Seq. To determine whether similar observations can be gained from newer scRNA-Seq technologies, with increased numbers of cells profiled and fewer genes detected with high sensitivity, we performed Drop-Seq on a mouse model of heart failure (myocardial infarction: MI). Notably, heart failure is known to result in large global gene expression changes associated with cell-death, cellular infiltration and injury induced cellular remodeling that make its comparison to controls more complex (Duan et al. 2017). This complexity was denoted in the joint-alignment of the sham and MI scRNA-Seq using Seurat version 2, in which new cell-populations are inappropriately predicted (endothelial), that are specific to the disease state (**Fig. S4B**). Applying cellHarmony to Seurat clustering of the Sham only (reference) finds that MI core-cellular identify gene expression is largely retained in all cell aligned populations, with more equivalent representation of cell types between datasets than with the joint-alignment (**Fig. 4A-C**). However, this data also suggests some multilineage gene expression priming within the fibroblast population, which adopts a weak combined smooth muscle, epicardial, and endothelial gene expression program, based on these default output heatmap and UMAP combined representations of the sham and MI data (**Fig. S4C**). To our surprise, the gene expression differences across all annotated cell populations (cell-type marker gene-based) indicate consistent gene upregulation within each specific cell-state, within the clustered cell states (regional), and within those shared between cell states (global) (**Fig. 4D**). However, while little gene downregulation is denoted in the combined heatmap representation of this data, we find regulation of diverse pathways with increased cell-state-specific gene expression. These changes are broadly reflected in comparable bulk RNA-Seq data from an independent study (Duan et al. 2017), indicating that differential expression is likely valid rather than technical (**Fig. 4E**). Moreover, examination of the predicted transcriptional regulatory networks for each specific cell-state comparison identifies well-defined MI-associated gene expression differences in human Fibroblasts (upregulation of *Col1a1, Col1a2, Col3a1, Col4a2, Col5a1, Col5a2* [Collagen scar] *Fn1, Thbs1, Mmp14*) along with novel implicated regulators which agree with prior literature (**Fig. 4F**) (van Dijk et al. 2008; Spinale et al. 2013; Frangogiannis 2017; Mouton et al. 2019). While the two central transcriptional nodes in this network, *Hif1a* and *Runx1*, have been previously implicated in MI and even as possible targets for therapy, these markers have not been explicitly associated with the MI fibroblasts (Kido et al. 2005; Semenza 2014; McCarroll et al. 2018). While another central node in this fibroblast network, *Egr1* was down-regulated, we note that in the Epicardial network, *Egr1* was also central but upregulated, in combination with other immediate-early central node genes *Jun* and *Fos* (**Fig. 4G**). This observation is important, as *Egr1* is a well-described transcriptional regulator and target for therapy in MI and is likely playing very different roles in these two different cell-types (Bhindi et al. 2006; Bhindi et al. 2012; Ramadas et al. 2014).

**Figure 4:**
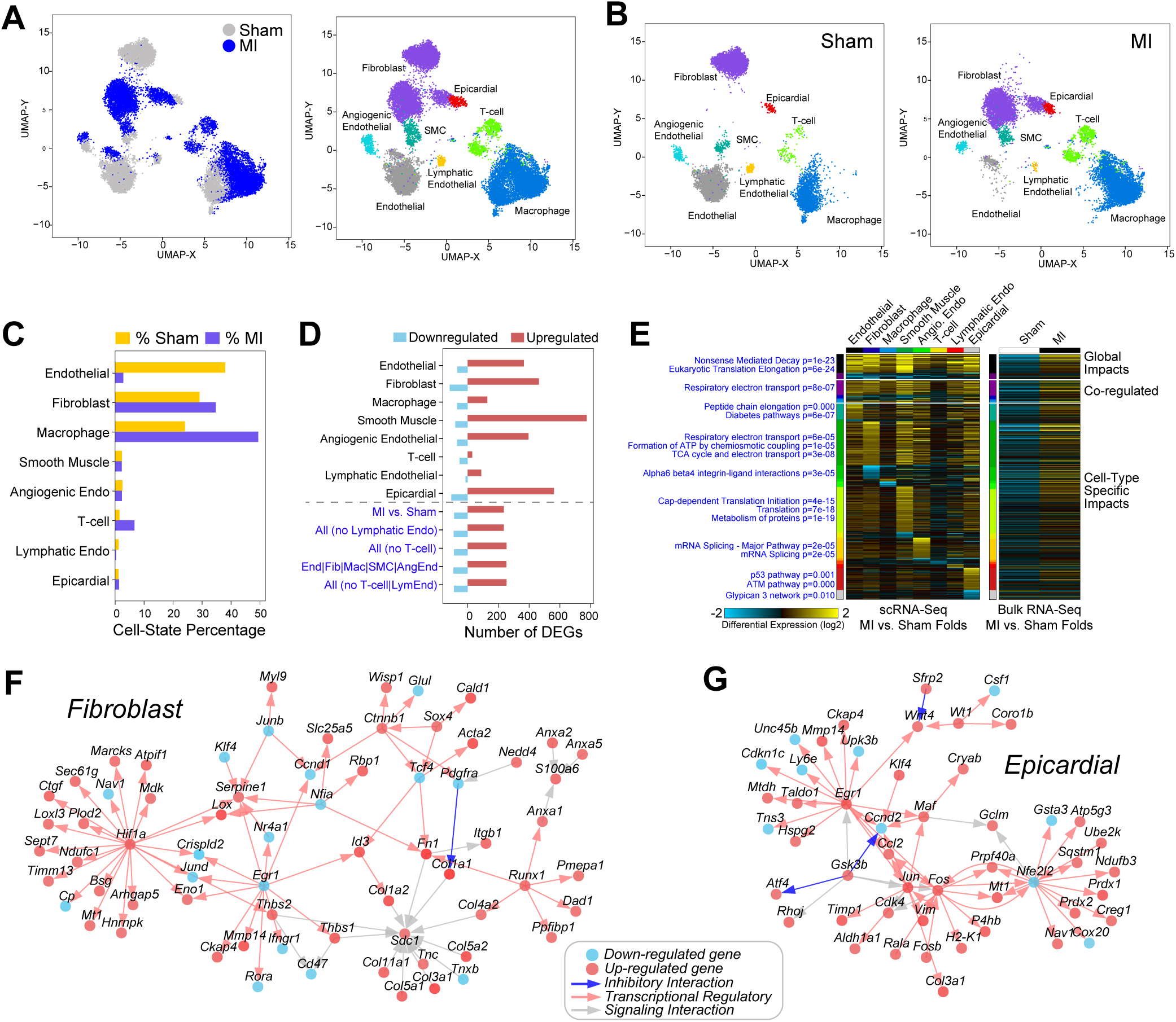
Identification of cell-type specific regulatory networks in heart failure. A) cellHarmony UMAP projection of mouse heart scRNA-Seq, comparing myocardial infarction (MI) to non-diseased hearts (Sham surgery). Cell cluster labels and colors were assigned by cellHarmony (MI) or from the Seurat analysis (Sham). B) Separation of the combined UMAP projections for reference Seurat Sham clusters and aligned MI clusters. Differences in the projected population locations in the MI are attributed to disease-associated transcriptomic changes. C) Cell-state population percentages for cells from Sham and MI. D) Number of differentially expressed genes for each cell-state (local), global (all AML cells versus all wild-type) and regional (coregulated cell-state clusters) comparisons for MI versus Sham (fold>1.5, empirical Bayes moderated t-test p<0.05 (FDR adjusted)). E) Ordered gene expression changes (upregulation and downregulation) for each global, regional and the cell-state specific comparison, based on relative gene expression p-value rankings, along with associated statistically enriched pathways and enrichment p-values (blue text) (left panel). The same genes are shown in the right panel for comparable bulk RNA-Seq of MI versus Sham, from a prior study (GSE96561). F,G) cellHarmony produced gene network displaying putative regulatory interactions (red arrows) among up-regulated (red notes) and down-regulated genes (blue nodes), separately for fibroblast (F) and epicardial (G) MI versus Sham comparisons.

### Using cellHarmony to identify diagnostic and prognostic gene networks

The evaluation of mouse disease data provides a well-controlled use case for determining valid differential expression changes, not confounded by genetic or clinical differences. To see whether cellHarmony is effective at identifying cell population-level genomic differences relevant in diseased patient samples, we analyzed human leukemia scRNA-Seq datasets before, during and/or following therapy. To minimize potential batch and donor effects for differential analyses, we selected scRNA-Seq from a pre-transplantation leukemia bone marrow biopsy relative to a post-transplantation biopsy on the same patient, although the donor genetics will differ from the recipient (Zheng et al. 2017). Alignment of the diagnostic (query) to the post-transplantation sample (reference) found an expected decrease in cellular diversity in the leukemia diagnostic sample, significant expansion of erythroblast compartment, and a global shift in the transcriptome of lymphoid, myeloid, and HSC assigned cells in the leukemia by UMAP visualization (**Fig. 5A-C** and **Fig. S5A**). These data suggest that this patient exhibits a wide-spread increase in the Erythroblast compartment prior to transplant (∼5 fold increase), with a broad decrease in early progenitor populations (e.g., HSC, megakaryocytic, erythroid, granulocytic, early-erythroid) and committed cell populations (e.g., naïve T-cell, platelets), compared to post-transplantation. Consistent with this observation, this patient was diagnosed with erythro-leukemia, which is characterized by proliferation of erythroblastic precursors (Zheng et al. 2017). Interestingly, the most divergent gene expression differences in pre-vs. post-transplant are found in Erythroblasts, which were characterized predominantly by gene down-regulation, where as other cell-types were principally characterized by up-regulation (**Fig. 5D**). Analysis of the predominant global and cell-state specific expression patterns reveal both significant cell-state-specific differences and shared global up-regulation in non-erythroblast cell populations (**Fig. 5E**). To determine whether such signatures were diagnostic for leukemia type, we analyzed bulk-RNA-Seq in a large un-annotated adult AML cohort consisting of 438 patients (Leucegene) and 16 donor CD34+CD45RA-cord-blood samples for the same gene modules (Lavallee et al. 2015). Using this individual erythro-leukemia patient, we identified two shared gene modules in over 120 Leucegene bone marrow biopsies, corresponding to a GATA1 transcription target cluster (AML-erythroblast-specific) and an inflammatory signature, indicated by the cellHarmony heatmap gene-set enrichment analysis (interferon-gamma and TLR). Comparison of patients with the high-GATA1 signature (and low inflammatory) to the high inflammatory (and low-GATA1) yields genes with expected enrichment in previously described erythro-leukemia and inflammatory/autoimmune disorder disease ontology terms, respectively, providing evidence that these signatures are indeed diagnostic (**Fig. 5F**).

**Figure 5.**
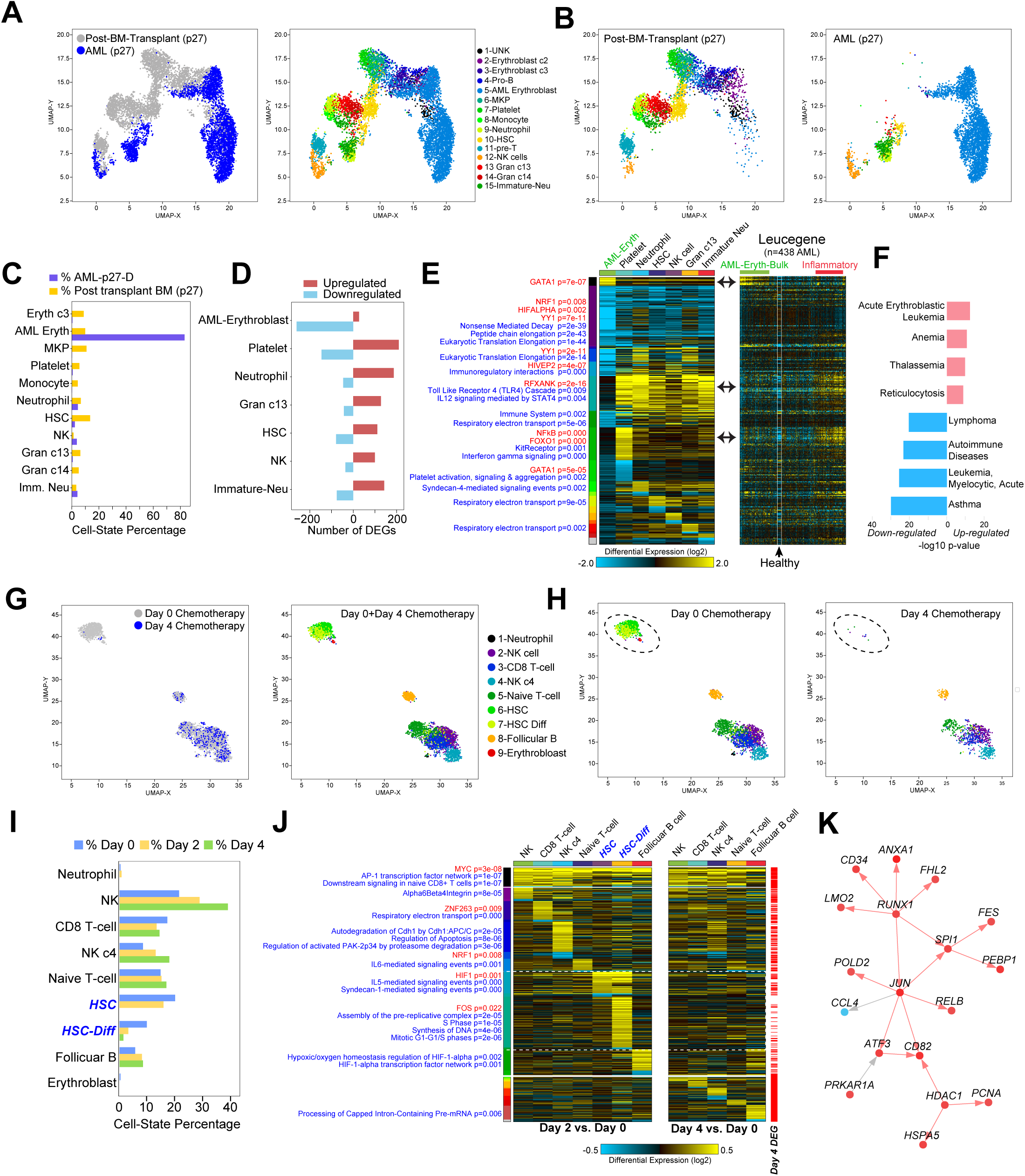
Assessing temporal changes in cell-state specific gene regulatory programs in Acute Myeloid Leukemia. A) UMAP projection of ICGS-NMF scRNA-Seq clusters from bone marrow mononuclear cells from a single-patient (AML027) after bone marrow transplantation and a cellHarmony aligned diagnosis AML sample. B) Separation of the combined UMAP projections for reference post-transplantation clusters and aligned AML clusters. C) Cell-state population percentages for cells from post-transplantation and AML. The diagnosis AML erythroblast cluster represents the dominant cell population in the AML patient. D) The number of corresponding gene expression changes in the AML compared to the post-transplantation. The AML erythroblast cluster is characterized by downregulation while all other clusters by upregulation. E) Ordered gene expression changes (upregulation and downregulation) for each global, regional and the cell-state specific comparison, with transcription factor target gene-sets displayed on the left (red) and pathways (blue) to the left of the heatmap with corresponding enrichment p-value (left panel). The same genes are shown in the right panel for comparable bulk RNA-Seq from the Leucegene AML RNA-Seq cohort with 438 patients and 16 donor CD34+CD45RA-cord-blood samples (row median normalized). Samples indicated as AML Eryth are those with a matching GATA1 transcription factor target enrichment in the left panel heatmap and those denoted as inflammatory correspond to cell states with an enrichment in interferon-gamma and TLR gene set enrichments. F) Disease Ontology gene-set enrichment with the program ToppGene of the two Leucegene patient groups high GATA1 and low inflammatory gene expression versus the converse sample sets. Red bars indicate statistical enrichment of upregulated (GATA1 high) gene sets and blue, downregulation associated genes (inflammatory). G,H) Analogous plots to those shown in panels A and B, respectively, for two scRNA-Seq timepoints of a single-patient’s blood at Day 0 and Day 4 of chemotherapy. I) Cell-state percentages for Day 2 and day 4 cells aligned to Day 0 ICGS-NMF clusters. No cells were aligned for HSC cells at Day 4 and only 11 cells for HSC-Differentiating at day 4 (too few for differential expression analysis of those clusters). J) Ordered AML time-course heatmap, in which ordered genes for both Day 2 and Day 4 versus Day 0 were combined. Differentials for Day 2 are shown on the left panel and Day 4 on the right, with red tick marks denoting genes significantly differentially expressed in Day 4 versus Day 0. K) The cellHarmony gene network for the differentiating HSC cluster at Day 2 versus Day 0.

As a final evaluation, we considered a recent AML treatment scRNA-Seq time-course (days 0, 2 and 4) with a combinatorial drug therapy to induce durable remission. The study authors use this data to demonstrate a loss in blood leukemic stem cell populations during therapy and metabolic disruption. Similarly, cellHarmony automatically indicates the same gradual loss in stem cell blood populations during treatment and enriched gene expression changes in analogous metabolic pathways (**Fig. 5G-I, Fig. S5B, Table S1**). However, our analysis also implicates time and cell-type dependent transcriptional regulatory networks, which suggest that the major population gene expression impacts happen at day 2 of therapy (global and cell-state specific upregulation) (**Fig. 5J**). Among the gene expression differences most pronounced at day 2, were those restricted to the two Leukemia stem cell-like populations. Examination of these changes reveals dozens of highly significant impacted pathways, notably syndecan-1, IL5, TGF-beta, EGFR, IFN-gamma, IGF-1, ErbB1, integrin and mTOR mediated signaling pathways (**Table S1**). These impacted pathways have well-described regulatory roles in proliferative signaling pathways, which is enriched among upregulated genes (Mitotic G1-G1/S phases) in concert (**Fig. 5J**). These differences were further exemplified by the cellHarmony predicted gene-regulatory network for differentiating HSCs, which highlight a core stem cell and proliferative regulatory program orchestrated by upregulation of *RUNX1, JUN* and *HDAC1* (**Fig. 5K**). Such transcriptional differences represent new potential biomarkers for intermediate response to therapy as well as novel possible therapeutic targets.

### Considerations for future applications of cellHarmony

While the application of cellHarmony to multiple real-world datasets demonstrates the power of this approach, as with any scRNA-Seq comparative dataset method, important caveats and validations should be employed by the user to ensure accuracy of the obtained alignment and differential expression results. First, batch and donor effects should be carefully considered and independent validations performed to assess the validity of the results. In our evaluations, these include bulk RNA-Seq analyses in which discrete cell population-level differences in both gene expression and cell frequency can be independently confirmed. Correcting for batch effects remains a challenge in a scRNA-Seq single-replicate studies. For this reason, cellHarmony includes a dedicated replicate dataset merging function, called cellHarmonyMerge, to integrate independent ICGS outputs from different reference samples to form a combined reference for use with cellHarmony alignment. When using cellHarmonyMerge, populations that are unique to only one of the references remain unique, while similar populations between the two references are combined into a single-cluster (**Fig. S2F**, Methods). As previously illustrated, use of this method can effectively side-step batch effects (Hay et al. 2018).

Second, we anticipate that cell populations may frequently be missing in the reference and present in the query or vice versa. When this happens, it is more likely for a novel cell population in the query to be inappropriately annotated as a specific reference cell-type, which shares partial transcriptomic similarity. For this reason, it is advised that users switch the query and reference in an additional analysis and examine the resulting cluster assignments. Such evaluations can assist in further adjustments to the default community alignment correlation cutoff to prevent incorrect associations. As demonstrated by our analyses, core-gene expression programs for cell-identity are retained in most disease states when compared to healthy (leukemia, heart failure) (**Fig. S4**). Such programs allow cellHarmony to effective project disease cells within a continuum of healthy transitional states, as illustrated here by an analysis of murine leukemia cells. To similarly obtain such biological insights, users are encouraged to develop annotated references with reasonably accurate cell-population labels to facilitate interpretation of cellHarmony’s findings.

Finally, when computing differences in the expression of populations, misinterpretation is possible due to too few cells aligning to a cell population or extreme data sparsity (e.g., low sequencing depth). In such cases, we recommend evaluating different thresholds for statistical significance in differential expression analyses and again using external data for validation (bulk-seq). Additional user recommendations on how to optimize parameters for each analysis are provided in the **Supplemental Information**.

## Discussion

Single-cell RNA-Seq continues to enable exciting insights from healthy and diseased tissues. Standardized and efficient workflows to explore such diversity benefit the research community by highlighting global and cell-state specific programs that can otherwise remain hidden. cellHarmony provides such reproducibility through the relatively fast alignment of independent scRNA-Seq graphs using a community alignment approach. As such, the approach remains scalable to large datasets, due to its low memory footprint, and will be less sensitive to outlier effects, through the use of neighborhood matching. While this method can accurately align diverse cells from different tissues and across different single-cell technologies, its principle power is in the determination of global, regional, and local differences among cell-states at the transcriptional, pathway, and gene-network level. As demonstrated, such insights include the localization of critical disease transcriptional changes to specific cell types, improved understanding of the specificity of drug targets to specific cell types in disease, improved diagnostic biomarkers, and novel regulatory and signaling networks that can inform therapy.

While cellHarmony is able to yield exciting new insights from disease scRNA-Seq datasets, a number of important challenges still remain. These challenges include the comparison of large single-cell patient cohorts, with variable cell numbers per sample, confounded by batch, genetics, and other effects and the explicit integrated analysis of multi-time-point datasets. Although new specialized and integrated approaches are needed to address these goals, our approach provides an important starting framework for more complex study designs involving scRNA-Seq through community alignment. We aim to address such needs in future iterations of this tool. We further note that development of methods for joint-alignment of scRNA-Seq datasets are continuing at an astonishing pace, which in some cases can eliminate donor and disease-biased clustering (e.g., Seurat version 3.0). While such approaches remain important in the integration of discrete datasets within a unified view, healthy reference single-cell maps will continue to provide an important ground-state truth for comparisons in which the determination and annotation of cell types is not conflated with disease or other secondary effects. With increasing use and decreasing expense of scRNA-Seq technologies, we anticipate approaches such as cellHarmony to become necessary to derive higher order insights into the investigation of pharmacological and disease heterogeneity.

## ACKNOWLEDGMENTS

We thank Harinder Singh for his critical discussions related to this method. This work was supported by Cincinnati Children’s Hospital Research Foundation and funding from the National Institutes of Health R01CA196658, R01HL122661, R21AI35595 (HLG) and R01CA226802 (NS).

## AUTHOR CONTRIBUTIONS

The manuscript was written by NS, HLG and ED. The method was conceived by NS and HLG. PD and NS implemented the algorithm. IVA and BCB designed and performed experiments. Computational analyses and algorithm evaluations were performed by ED, DS, KF, SH and NS.

## DECLARATIONS

All software is open-source and provided in Github according to the Apache License 2.0. Associated datasets are published and/or provided as example datasets with the accompanying software (DemoData/cellHarmony directory).

## SUPPLEMENTAL FIGURE LEGENDS

**Figure S1. Using cellHarmony to minimize clustering bias.** A) T-SNE plot of the joint-alignment of two human bone marrow scRNA-Seq samples from the Human Cell Atlas initiative (Hay et al. 2018) using the software Seurat canonical correlation analysis (CCA) (Seurat version 2.3.4). The panel of the left panel indicates CCA clusters, the middle indicates donor source, and the right indicates cell-type annotations provided by Hay et al. B) Donor-related biases in clustering are denoted by the percentage of cells associated with the two donors in all clusters (ranked from most impacted to least). C) Heatmap displaying the cellHarmony alignment of donor 5 (BM5) to the reference ICGS-NMF clusters of donor 2 (BM2). D) Donor-related biases in alignment of BM5 to BM2 cells, as shown in panel B, for cellHarmony. Note, BM2 and BM5 were noted by Hay et al. to have different cell cluster frequencies to but that shared cells for all reported clusters. E) Comparison of donor biased clusters in cellHarmony and Seurat. The Y-axis denotes the number of clusters with “high” donor bias, and the x-axis indicates the relative percentage of cells in each cluster differing between donors. This percentage is calculated as the absolute difference in the percentage of cells for each donor, divided by the maximum percentage: abs(%BM2 - %BM5)/max(%BM2,%BM5).

**Figure S2. Alignment of previously defined multi-tissue mouse cell atlas clusters with cellHarmony.**

A) cellHarmony alignment results as a combined heatmap for a large mouse cell atlas (Tabula Muris), corresponding to >47,000 10x Genomics scRNA-Seq profiled cells from 12 tissues (query) against >6,000 annotated SMART-Seq2 scRNA-Seq profiles from the same tissues. Cells in both datasets were restricted to those which had common Cell Ontology labels across technologies, within each cell-type (e.g., Lung Macrophage labeled from SMART-Seq2 and Lung Macrophage labeled from 10x Genomics). Gene clusters were defined based on the Cell Ontology labels from the SMART-Seq2 dataset, with markers derived using the MarkerFinder function within AltAnalyze (top 60 reported markers for each annotated tissue by cell-type). At the top of the heatmap are clusters denoted by different colors and on left are genes associated with those clusters. Although some cell-types (e.g., B-cells) will have near identical transcriptomes in different cell-types, these were still kept as independent clusters for alignment, but will not always have unique marker gene clusters. Cells corresponding to the two different technologies are displayed below the heatmap (not all white and black lines will be visible). B) Percentage of cells for each cell-population that were defined by the original study authors (SMART-Seq2) or by cellHarmony alignment (10x Genomics). Note, for each technology, different isolation methods were employed (FACs in SMART-Seq2 and unbiased in 10x Genomics). C) Classification of the 47k 10X Genomics dataset against SMART-Seq2 reference centroids (clusters and marker genes derived from panel A using the software Reference Component Analysis (RCA). Heatmap output of RCA, showing correlation of 46 centroid references (columns) and all single-cells (rows), with “ground truth” cell classifications for each cell indicated on the column color bar. The calculated ARI is for agreement between the 46 Cell Ontology clusters and RCA de novo clusters. D) Comparison of cell alignment approach Adjusted Rand Index agreement values for the different evaluated methods, across different filtering thresholds (Pearson correlation cutoff) for the 47k 10x Genomics query against the 6k SMART-Seq2 reference (46 Cell Ontology groups). Note, scmap does not report alignment scores directly, hence, a single reported value is shown. cellHarmony using either ICGS-NMF reference profiles or using the profiles from panel A. E) Specificity of alignments from 10x Genomics cells (50% test set) against 10x Genomics cells (50% training set) for the original author denoted Cell Ontology terms. Increased saturation of the squares indicates a higher proportion of assignments, with the darkest green representing 100% assignment in that category. Note, B-cell, T-cell and leukocyte ambiguity across tissues, as expected. F) Histogram of cellHarmony alignment Pearson correlation coefficient values for aligned cells in the 10x Genomics 50/50% test and training set (grey bars), with all cells included versus exclusion of kidney collecting duct (red bars) or liver hepatocyte (blue bars) profiles from the reference. Both kidney and liver cells should not have an analogous reference in the filtered training reference set, hence, alignments should be poor.

**Figure S3. Alignment of de novo multi-tissue mousce cell atlas clusters with cellHarmony.** A) Similar cellHarmony alignment heatmap as shown in Fig. S2A but using de novo clusters derived from the independent ICGS-NMF analysis of all 12 10x Genomics tissues for the 50/50% test and training datasets. The 171 clusters represent transcriptionally unique cell populations which can include cells from multiple tissues in a single cluster based on the cellHarmonyMerge function. B) The Adjusted Rand Index for the results displayed in panel A using cellHarmony 171 reference cluster profiles (MF=MarkerFinder), scmap or centroid-based classification in cellHarmony (171 cluster centroid). Confidence intervals for ARI were computed using the normal approximation.

**Figure S4. Evaluation of differential expression analysis algorithms and disease-reference alignments**. A) To assess the suitability of different algorithms for scRNA-Seq differential expression relative to bulk RNA-Seq, Venn diagrams are shown comparing reported differentially expressed genes (DEGs) from T-cells and B-cells profiled either by bulk RNA-Seq or scRNA-Seq (**Supplemental Information**). Three algorithms for differential gene expression analysis (eBayes, SCDE and MAST) were applied for scRNA-Seq, compared to “gold-standard” results from bulk-RNA-Seq, demonstrating the relative effectiveness of the eBayes method. B) Joint-alignment analysis results (Seurat CCA version 2.3.4) for healthy (Sham) and diseased (myocardial infarction: MI) viewed as a t-SNE plot for the reported clusters (left) and samples (right). Sample biased clusters in the t-SNE plot are labeled according to the annotated cell-types (marker defined). C) cellHarmony combined heatmap of the MI (query) and Sham (reference) alignments, with cell-type labels derived from AltAnalyze gene-set enrichment analysis and previously described marker genes, for the obtained Sham Seurat clusters.

**Figure S5. cellHarmony comparison of acute myeloid leukemia patient samples overtime time.** A) cellHarmony heatmap human bone marrow mononuclear cells (BMMCs) from an AML patient at diagnosis (query) aligned to a post-transplantation biopsy (reference) from the same individual. Cell-type predictions are using input gene-sets from Hay et al. with AltAnalyze (derived human bone marrow cell population markers). Reference AML diagnosis cell populations were determined using ICGS-NMF. B) Similar heatmaps as in A, with peripheral blood cells from AML, 2 days (left) or 4 days (right) after the induction of chemotherapy (query) relative to Day 0 cells (reference). Reference AML Day 0 cell populations were determined using ICGS-NMF.

