## Supplemental-Information for "cellHarmony: Cell-level matching and holistic comparison of single-cell transcriptomes"

SUPPLEMENTARY INFORMATION

**User and Command-Line Interfaces**

cellHarmony can be run through the AltAnalyze graphical user interface or through the command-line. AltAnalyze is available as pre-compiled binaries (http://www.altanalyze.org) or as source code (PyPI python 2.7 installation or https://github.com/nsalomonis/altanalyze). Both command-line and pre-compiled binaries can be run on the command-line (https://github.com/nsalomonis/altanalyze/wiki/CommandLineMode). The cellHarmony-Align code can also be independently run as an alternative to AltAnalyze (https://github.com/AltAnalyze/cellHarmony-Align). This code-base only performs the community-alignment function, without the graphical outputs (heatmap, UMAP, networks) or differential expression. When run through AltAnalyze from the graphical user interface, the user must first install a species database from the command-line, for example:

```
$ python AltAnalyze.py --species Hs --update Official --version EnsMart72 --additional all
```

Or from the graphical user interface from the main menu (Add New Species button) or when prompted when first started (https://altanalyze.readthedocs.io/en/latest/RunningAltAnalyze/). To open the cellHarmony user interface, start AltAnalyze, proceed to the main menu, select the appropriate downloaded species option, proceed to the Additional Analyses menu and the Cell Classification menu to begin the analysis. A description of the input files for analysis are described in the following section.

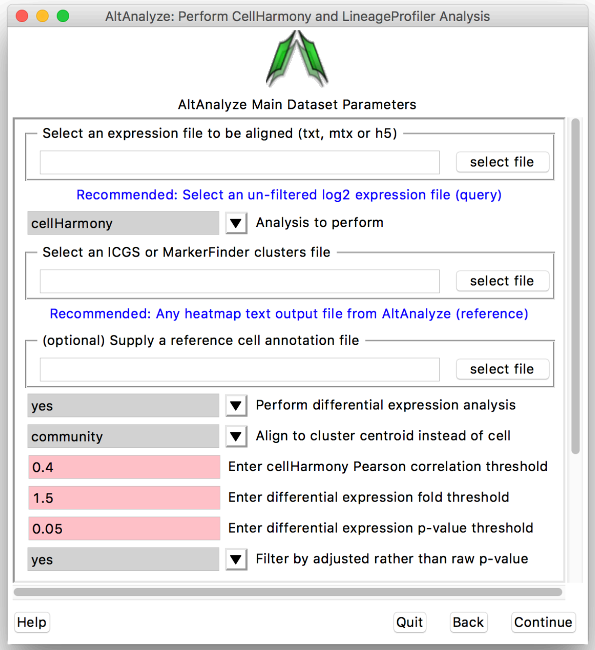

For the command-line interface, cellHarmony can be run as a single command:

```
$ python AltAnalyze.py --cellHarmony yes --input queryFolder/AML.txt --reference
referenceFolder/ICGS-NMF/FinalMarkerHeatmap.txt --platform RNASeq --species Mm --
correlationCutoff 0.4 --referenceType community --performDiffExp True --cellHarmony
yes --adjp True --fold 1.5 --label /labels/BoneMarrow_cluster_names.txt
```

When only performing alignment:

```
$ python AltAnalyze.py --cellHarmony yes --input queryFolder/AML.h5 --reference
referenceFolder/wild-type.h5 --platform RNASeq --species Mm --correlationCutoff 0.4 --
referenceType community --performDiffExp False --cellHarmony yes --label
/labels/BoneMarrow_cluster_names.txt --genes markerGeneFolder/markerGenes.txt
```

**Running cellHarmony on a Cluster**

We have observed that cellHarmony may produce an error when run on a cluster, specifically when nodes without Advanced Vector Extensions (AVX or AVX2) are selected. This is due to a requirement of the community clustering python libraries. To avoid this issue, request a node with AVX. For example, using bsub: bsub -M8000 -W 4:00 -J testname -n 4 -R "span[hosts=1]" **-R avx, avx2** -o log.out

**Input Data Files**

cellHarmony can be run in three major modes: 1) "rapid-mode" of two scRNA-Seq datasets, 2) alignment and visualization with no differential expression, and 3) full analysis workflow. Options 2 and 3 require an AltAnalyze format heatmap text file for data visualization and marker gene selection. This heatmap includes the output of unsupervised population detection with ICGS version 2 (ICGS guide-gene folder, ICGS-NMF markers folder or DataPlots/Marker finder output or other produced heatmap). Instructions for creating this file from the output of Seurat can be found on the cellHarmony website (http://www.altanalyze.org/cellHarmony/). When only aligning two scRNA-Seq files (rapid-mode), users must supply the following input:

- (File option 1) Two sparse-matrix (h5 or mtx format) scRNA-Seq dataset files (query and reference). For mtx files (Cell Ranger, version 1-3), the mtx directory must contain a barcodes.tsv, genes.tsv, or features.tsv file. cellHarmony currently supports only h5 or mtx files from the same version of Cell Ranger.

- (File option 2) Two tab-delimited text files with log2 read-normalized expression data (query and reference). These files can contain all genes or be restricted only contain marker genes if desired. The file must contain a gene column (column 1) and a barcode row (row 1).
- Tab-delimited marker genes text file. Genes in column 1.
- (Optional) A tab-delimited label file, with cell barcodes/IDs (column 1) and labels (column 2).

The output of the rapid-mode analysis is a tab-delimited text file with: A) the query and aligned reference cell barcodes, B) aligned communities (cell partitions) and C) user-supplied reference labels (optional). In addition to these same input files, to obtain the other standard outputs of cellHarmony, users must additionally supply:

- An ICGS results directory heatmap file (see above directory locations) along with an ExpressionInput folder in the same parent directory containing the full gene normalized expression file (log2 or non-log). For Seurat, users must save the equivalent format files in the same folder structure. A script is provided at: http://www.altanalyze.org/cellHarmony/ to produce this output from R. This heatmap text file should be selected as the cellHarmony reference.

The query file can be a tab-delimited or sparse matrix input. If sparse, this file will be exported to a dense format, column normalized log2 file to the same directory as the h5 or mtx file (options 2 and 3). This file will be used for joint visualization (heatmap, UMAP) of the query and reference and downstream differential expression, network, pathway, and systems analyses.

Batch effects should be evaluated, and if necessary, corrected prior to cellHarmony alignment using external tools (e.g., MNN). Example reference and query datasets and instructions are provided with the AltAnalyze software (DemoData) and online (https://github.com/AltAnalyze/cellHarmony-Align/tree/master/sample_data).

**Combining ICGS Datasets**

As an optional reference, cellHarmony can merge multiple ICGS outputs for diverse datasets (cellHarmonyMerge function). This method is recommended for cases in which users want to combine and compare multiple scRNA-Seq datasets simultaneously. The combined analysis uses the union of all supplied marker genes, averaged similar-cell profiles, and a separate file with all combined and ordered cells. To produce these results, the cellHarmonyMerge function: 1) selects all unique marker genes from the collectively provided set of inputs, 2) imports the expression data for these genes from the original complete gene matrix (e.g., AltAnalyze ExpressionInput "exp." file) and converts these to log2 values, 3) computes centroid expression for cells within the same identified cluster for all marker genes, 4) averages similar centroids "clusters" based on all pairwise centroid comparisons (Pearson correlation>0.9, default) to produce merged centroids, 5) filters the combined dataset to only include genes with non-zero values for all columns, 6) re-clusters the new reference centroids (HOPACH clustering) to produce a re-ordered matrix of reference cell-population centroids, 7) identifies and ranks marker genes for each of the combined clusters (MarkerFinder function, AltAnalyze) and 8) outputs a cell-level output for cellular barcodes corresponding to the cluster centroids for the final marker genes. The resultant all-cells reference text file can be used for downstream cellHarmony analyses. As an example, we have applied this workflow to 35 distinct cell populations we identified from bone marrow RNA-Seq generated by the Human Cell Atlas project, following independent ICGS analyses of each independent donor (Hay et al. 2018).

**Programmatic Options and Parameter Tuning**

When cellHarmony is run from the command-line, it requires specifying the following the user defined options: 1) the cell-alignment correlation cutoff (rho>0.4, default), 2) variable marker genes, 3) differential expression p-value threshold (pval<0.05, default), 4) differential expression fold-change threshold (fold>1.5, default) and 5) FDR adjustment (adjp=True, default).

The code can be run as an independent python script in an alignment-only “rapid-mode” (https://github.com/AltAnalyze/cellHarmony-Align) or through the full cellHarmony programmatic or graphical user interface. When run in “rapid-mode”, the user can modify multiple parameters for community clustering including the number of nearest neighbors (default =10) and number of trees for Annoy (default = 100), at which level to cut what level to cut the Louvain clustering dendrogram (default = 0), the minimum correlation for reporting a matching cell (default = -1), and the genes to consider for analysis. To optimize speed and memory utilization, h5 files are recommended. On a Mac OSX laptop with 16GB of RAM (2.9 GHz Intel Core i7), alignment of ~2,100 cell query cells to a ~3,800 cell reference completes in under a minute (https://github.com/AltAnalyze/cellHarmony-Align/tree/master/sample_data). With samples with weak heterogeneity (few community clusters) but many cells, the analysis time can be slower, (~10,000 cell query to a ~13,000 cell reference completing in 29 minutes with ~325MB maximum RAM used, with only 31 and 43 communities detected). Other alignment options present in the software include all cell-by-cell or cell-by-centroid alignments using k-nearest neighbor classification (KNN)(K=1), although community clustering is recommended (default). When applying these alternative comparisons, the nearest neighbor of a query cell profile is assigned to its best match in the reference set based on all possible cell-cell Pearson correlations.

The major determinants of which options to use are the experimental platforms compared (e.g., Fluidigm versus sci-CAR), increased or decreased data sparsity, and poorly detected heterogeneity within clusters. We recommend these parameters be tuned based on the following criteria: A) decreased or no fold-change threshold for highly sparse transcriptional data with excessive dropouts, B) use of non-adjusted moderated t-test for highly sparse data, C) use of an alternative ICGS/Seurat reference for highly sparse dataset with weakly detected heterogeneity, D) decreased correlation threshold when comparing highly dissimilar scRNA-Seq protocols/platforms, and E) increased correlation threshold in datasets where contaminating cells/doublets are present in the query or reference. No clear difference has been observed in our benchmarking of different ICGS clustering outputs (Euclidean versus Cosine) or use of marker genes in *a priori* defined clusters (see Mouse Cell Atlas example). Furthermore, if questionable alignments occur, the user is recommended to reverse the query and reference and re-run cellHarmony to determine whether cell or population predictions significantly differ (requires ICGS analysis of the new reference).

**Output Files**

As output, cellHarmony produces multiple tabular and visualization results, depending on the type of reference dataset supplied (full expression matrix versus pre-filtered). The initial outputs of cellHarmony are: 1) final association z-score matrix derived from the Pearson correlation coefficients for all cells (labels optional), 2) expression matrix in which each cell is placed adjacent to its best match, 3) query-only cell matrix with cells ordered and annotated according to the classification, 4) gene expression heatmaps of the expression matrices, 5) cell frequency and gene expression difference bar charts, 6) statistical differences in the frequency of aligned cell populations between reference and query samples, 7) statistically significant genes for all comparisons and associated values, 8) UMAP projection of the query and reference cells combined, 9) Pattern ordered heatmap with of fold differences in all compared cell populations (global, regional, local) with enriched Pathway Commons or transcription factor binding site gene-sets, and 10) network plots and gene interaction lists. The log2-normalized expression profiles for both the reference and query are displayed as a combined heatmap, in the reference gene and cell order, with the query inserted alongside each cellHarmony match to assess their relative similarity. The frequency of cells present or absent from the query in each cell population is further reported and statistically quantified using a Fisher exact test to allow for the assessment of the lineage impact with cellular, molecular, or genetic perturbation in the query.

**Evaluation Datasets**

The following datasets were selected for evaluation within cellHarmony as they contained defined populations or have well-described cellular biology. Associated analysis scripts, input data files, and results can be obtained at https://www.synapse.org/#!Synapse:syn18500191.

Tabula Muris Mouse Cell Atlas (MCA): Gene cell matrices for all SMART-Seq2 and 10x Genomics datasets were downloaded from GEO (GSE109774) and processed in ICGS version 2 using the software default options. Tissues samples with replicates were combined following counts normalization and log2 adjustment in AltAnalyze (SMART-Seq2 = CountsNormalize.py, 10x Genomics = ChromiumProcessing.py script) and jointly analyzed (MergeFiles.py script) (https://github.com/nsalomonis/altanalyze). Annotations for each MCA cell were obtained from the study authors (https://github.com/czbiohub/tabula-muris/tree/master/00_data_ingest/03_tissue_annotation_csv). ICGS cell clusters were joined using the cellHarmonyMerge function, with correlation merge threshold=0.95 and centroid=True, as an optional reference for cellHarmony testing with high-resolution cluster reference evaluation, which identified 171 non-redundant clusters from the 12 evaluated 10x Genomics dataset tissues (see above Synapse repository). cellHarmonyMerge markers from the automated MarkerFinder function were identified from this merged dataset for the training cohort (n=24,618 cell barcodes) to apply the test cohort (n=24,618 cell barcodes). For comparative algorithm evaluation, no gene filtering was applied with the exception of RCA, which required a pre-determined list of marker genes and pseudo-bulk centroids per defined cell cluster.

Bone Marrow Human Cell Atlas (HCA): Two samples were selected for scRNA-Seq comparison which previously displayed divergent cell population frequencies (BM2=50 yrs. old male, BM5=29 yrs. old male) (Hay et al. 2018). For each sample, all 8 independent captures were combined as previously described (Hay et al. 2018). ICGS version 2 was performed on BM2 using the default options and cell-types annotated using the bone marrow cell type markers (Hay et al. Table S5).

Transitional cell-states in bone marrow progenitor singlets: A dataset comprised of 383 hematopoietic bone marrow progenitor cells with high-confidence assigned cell-types and singlet-restricted profiles, validated via microfluidics cell capture imaging, was obtained from the GEO database along with the published ICGS unsupervised clustering results (GSE70245). Associated AML scRNA-Seq and bulk RNA-Seq were obtained from (GSE77849) and processed using the software RSEM to match the scRNA-Seq.

Murine Myocardial Infarction (MI): A previously unpublished scRNA-Seq dataset from a mouse model of myocardial infarction and sham surgery was produced for evaluation of cell-state specific transcriptomic differences (see Single-Cell RNA-Sequencing). This sequencing data, expression files, and metadata have been deposited in the open-access Synapse database (https://www.synapse.org/#!Synapse:syn18516494/files/). Seurat Canonical Correlation analysis (version

2.3.4) was used to demonstrate alignment biases (Sham and MI) and separate Seurat sample processing of each sample to compare by cellHarmony. Count matrices were filtered to cells having a minimum of 200 genes expressed, proportion.mito < 0.25 and >= 400 UMIs (Sham n=13,858; MI n=11,240). Standard Seurat processing was conducted, including log-normalization, regressing out nUMI, mitochondrial proportion and cell cycle indicators (proportion of histone and Seurat G2/M transcripts), and scaling. Canonical correlation vectors were computed (RunCCA) using the union of the top 20% of Seurat-determined variable genes in each sample and which were expressed in both samples. Based on diagnostic plots, sample alignment (AlignSubspace) was performed using the top 20 CCA dimensions. The Seurat FindClusters function with 20 CCA dimensions and resolution=0.8 identified 16 clusters. Marker genes for each cluster (FindConservedMarkers) were found and cell types were determined by comparisons to previously identified marker. The Seurat cellHarmony processing script (http://altanalyze.org/cellHarmony/) was used to convert the expression matrix, cell-state marker genes and cellular barcode clusters to an AltAnalyze compatible heatmap. Prior to cellHarmony analysis, predicted cell doublet profiles were excluded from the analysis after selecting the union of doublet predictions from the softwares DoubletDecon and Scrublet (Wolock et al. ; DePasquale et al. 2018). Corresponding bulk RNA-Seq data was obtained from GEO (GSE96561) and processed using the software Kallisto, called from within AltAnalyze.

Human Acute Myeloid Leukemia Datasets. Data from two separate patient-matched AML studies were analyzed. The first study was a diagnostic and relapse sample from a patient with erythro-leukemia (p27) and post-transplantation biopsy profiled using the 10x Genomics Chromium platform (version 1 chemistry), obtained from the 10x Genomics website (https://support.10xgenomics.com/single-cell-gene-expression/datasets/1.1.0/aml027_pre_transplant, https://support.10xgenomics.com/single-cell-gene-expression/datasets/1.1.0/aml027_post_transplant). The data were processed from the supplied sparse-matrix input files in AltAnalyze using ICGS version 2 to obtain the post-transplant ICGS-NMF heatmap text file as the reference for cellHarmony. A second AML scRNA-Seq time-course was obtained from GEO (GSE116481) as a combined gene-counts file and pre-processed using the AltAnalyze CountsNormalize.py python script. The day 0 diagnostic blood sample was used as the reference for cellHarmony following ICGS version 2 analysis. To identify less restricted gene expression differences at Day 2 and Day 4 compared to Day 0, cellHarmony was run without performing a p-value FDR adjustment and fold > 1.5. As a bulk RNA-Seq comparator, raw sequencing data from the Leucegene AML consortium (GSE49642, GSE52656, GSE62190, GSE67040) were downloaded and pseudoaligned to the Ensembl 72 transcriptome using Kallisto. These samples were combined with healthy CD34+CD45RA- cord-blood samples from the same laboratory (GSE48846).

**Single-Cell RNA-Sequencing**

Acute myocardial infarction (MI) was performed on C57BL6/J wild type (WT) 8-10-week-old male mice (The Jackson Laboratories and confirmed via echocardiographic analysis, similar to previously described (Duan et al. 2017). A left thoracotomy was performed via the fourth intercostal space and the lungs retracted to expose the heart. After opening the pericardium, the left anterior descending coronary artery was ligated with 7-0 silk suture approximately 2 mm below the edge of the left atrial appendage. Ligation was considered successful when the anterior wall of the left ventricle turned pale. The lungs were inflated by increasing positive end-expiratory pressure and the thoracotomy site closed in layers with 6-0 suture. Animals were maintained on a 37 °C heating pad until recovery and for 2 h after surgery. Another group of mice underwent sham ligation, with a similar surgical procedure without tightening the suture around the coronary artery. Mice with an estimated pressure gradient across the aortic constriction below 40 mmHg were not included in the experiments. Hearts were collected at 14 days post-Sham surgery (n=4, pooled) or MI (n=1), perfused with ice-cold PBS to remove red blood cells followed by perfusion with 50 mM KCl to arrest the heart in diastole and then fixed for 4 hours in freshly prepared 4% PFA at 4 °C, rinsed with PBS and cryoprotected in 30% sucrose/PBS overnight before embedding in OCT (Tissue-Tek). DropSeq was performed as previously described (Macosko et al. 2015). The quantity and quality of cDNA was measured using an Agilent Bioanalyzer hsDNA chip. To generate a library cDNA was fragmented and amplified (12 cycles) using the Nextera XT DNA Sample prep kit with three separate reactions of 600, 1,200 and 1,800 pg input cDNA. The libraries were pooled and purified twice using 0.7X volume of SPRIselect beads. The purified libraries were quantified using an hsDNA chip and were sequenced on an Illumina HiSeq 2500 using the sequencing parameters described in the DropSeq protocol. Reads were aligned to the mm10 mouse genome using Bowtie2 (Langmead et al. 2009) and tagged with the gene name of the overlapped exon. Gene reads were counted by unique UMIs per cell and a digital expression matrix was created. All animal procedures were performed and approved according to the Department of Laboratory Animal Medicine and the University Committee on Animal Resources at Cincinnati Children's Hospital Medical Center. scRNA-Seq data was deposited in Synapse (https://www.synapse.org/#!Synapse:syn18516494/files/).

**Evaluation Details for cellHarmony**

<u>cellHarmony Evaluation Parameters</u>: ICGS was run using AltAnalyze version 2.1.2 from input read counts or normalized count matrices using the software default options (Olsson et al. 2016). The selected references for cellHarmony were produced and tested: R1) ICGS version 2.0 primary output file (ICGS-NMF/FinalMarkerHeatmap_all.txt), R2) intermediate ICGS output (ICGS/Guide-3 result), R3) marker

profiles from previously derived cell clusters (supervised MarkerFinder analysis), or R4) the cellHarmonyMerge MarkerFinder output. ICGS outputs were produced using the default software options. Seurat-CCA was run as described above for the heart failure model analysis and HCA bone marrow progenitor dataset. For the Tabula Muris evaluation, gene-sets were tested for R1 and R3 (10x Genomics vs. SMART-Seq2) and R4 (10x Genomics test vs. training). For all Tabula Muris evaluations, no Pearson correlation cutoff was applied (rho>-1), since cells were restricted to those with common labels in both the SMART-Seq2 and 10x Genomics datasets (46 Cell Ontology labels). No Pearson correlation cutoff was used for the mouse AML or HCA bone marrow analyses, as these cells were previously determined to have mutually overlapping cell populations (Meyer et al. 2016; Hay et al. 2018). For all other analyses, the default Pearson correlation threshold was applied. For differential expression analyses, the default options were applied with the exception of the mouse AML and human AML sample p27 comparisons, in which a more stringent fold change was applied (2-fold change), as these datasets had few cells overall and few cells aligned to most cell populations. A non-adjusted moderated t-test was applied for the venetoclax treatment patient data to increase the detection of common gene expression differences during treatment.

Benchmarking Differential Expression Statistical Methods:

Differential gene expression estimates from the software cellHarmony (empirical Bayes), SCDE and MAST were compared using bulk RNA-Seq as a control. Bulk RNA-Seq T-cells and B-cells (GSE51984) were aligned to the human genome (hg19) with the program STAR and analyzed using AltAnalyze to identify differentially expressed genes (DEGs) with an FDR corrected $p < 0.05$. Single-cell RNA-Seq from human peripheral blood mononuclear cells (PBMCs) was downloaded from the 10x Genomics website (https://support.10xgenomics.com/single-cell-vdj/datasets) and processed in AltAnalyze with the ICGS algorithm to identify a CD8+ T-cell population and B-cell population. These data were compared in SCDE, MAST and AltAnalyze. For genes identified in the bulk and single-cell comparisons with a fold change in the same direction, DEGs were compared to calculate sensitivity and specificity.

**Evaluation Details for Evaluated External Algorithms:**

Reference Component Analysis (RCA): RCA associates individual cells with curated reference gene expression profiles through a correlation analysis., followed by a built-in method to define cell clusters (Li et al. 2017). As RCA does not come with the inherent capability to include external references, only the three references utilized in their manuscript, modifications to the code were required to compare the algorithm performance to that of cellHarmony and other alignment methods. Therefore, the following modifications were made to the RCA source code to allow for the addition of external references and the export of values required for assessment of accuracy:

1. The "sysdata.rda" file was obtained from https://github.com/GIS-SP-Group/RCA and loaded into R Studio (version 1.1.383, R version 3.4.2).
2. A new external reference, consisting of the described centroids and MarkerFinder marker genes, was inserted as the third reference dataset in the GlobalPanel list. This new "sysdata.rda" file was used for analysis following the vignette provided in RCA.
3. Modifications to the code to allow for use of this third reference without altering the core algorithm of RCA were made, with documentation of these modifications (https://www.synapse.org/#!Synapse:syn18500192).
4. Cell and gene names were reformatted to fit RCA standards, with these new files located at the Synapse link above.
5. In the final visualization step of RCA (RCAplot() function), column color labels were extracted from both the RCA_clusters and external_labels heatmaps, along with the corresponding cell IDs, to calculate Adjusted Rand Index (ARI).

As a surrogate for alignment agreement, centroids were calculated for the 46 cell populations in the Mouse Cell Atlas dataset (SMART-Seq2) and incorporated into the "sysdata.rda" file, as outlined above, with the 47k 10x Genomics test data input for RCA. For this assessment, maximum agreement is defined as ARI of 1, or perfect equivalency between the RCA defined cell clusters and the biologically validated "ground truth" cell states. Cluster definitions were derived from the RCAplot() heatmap visualization code, with the "ground truth" cell populations defined as the cluster labels in the heatmap with external labels and the RCA derived populations defined as the labels in the RCA clusters heatmap. ARI can be calculated for the results of RCA, but not accuracy, as the method does not allow for direct label projection.

<u>scmap</u>: scmap applies label projection from a reference dataset to a test dataset, by considering the agreement between multiple cell-similarity metrics. scmap was applied to two separate evaluation datasets: 1) 10x Genomics (47k) query cells compared to SMART-Seq2 (6k) reference and 2) 10x Genomics (24k test) query and 10x Genomics (24k training) reference. Both the query and reference were loaded as a "SingleCellExperiment" object which the scmap library uses as a scaffold. The "selected features" option was used to restrict the analysis to 500 variable genes for each dataset, based on expression and dropout-based rate. Clustering was performed using the "indexCluster" method, and projection was carried out using the standard scmap mapping algorithm, as outlined in the vignettes.
