## Supplementary figures and images for "cellHarmony: Cell-level matching and holistic comparison of single-cell transcriptomes"

### Supplemental-Figures

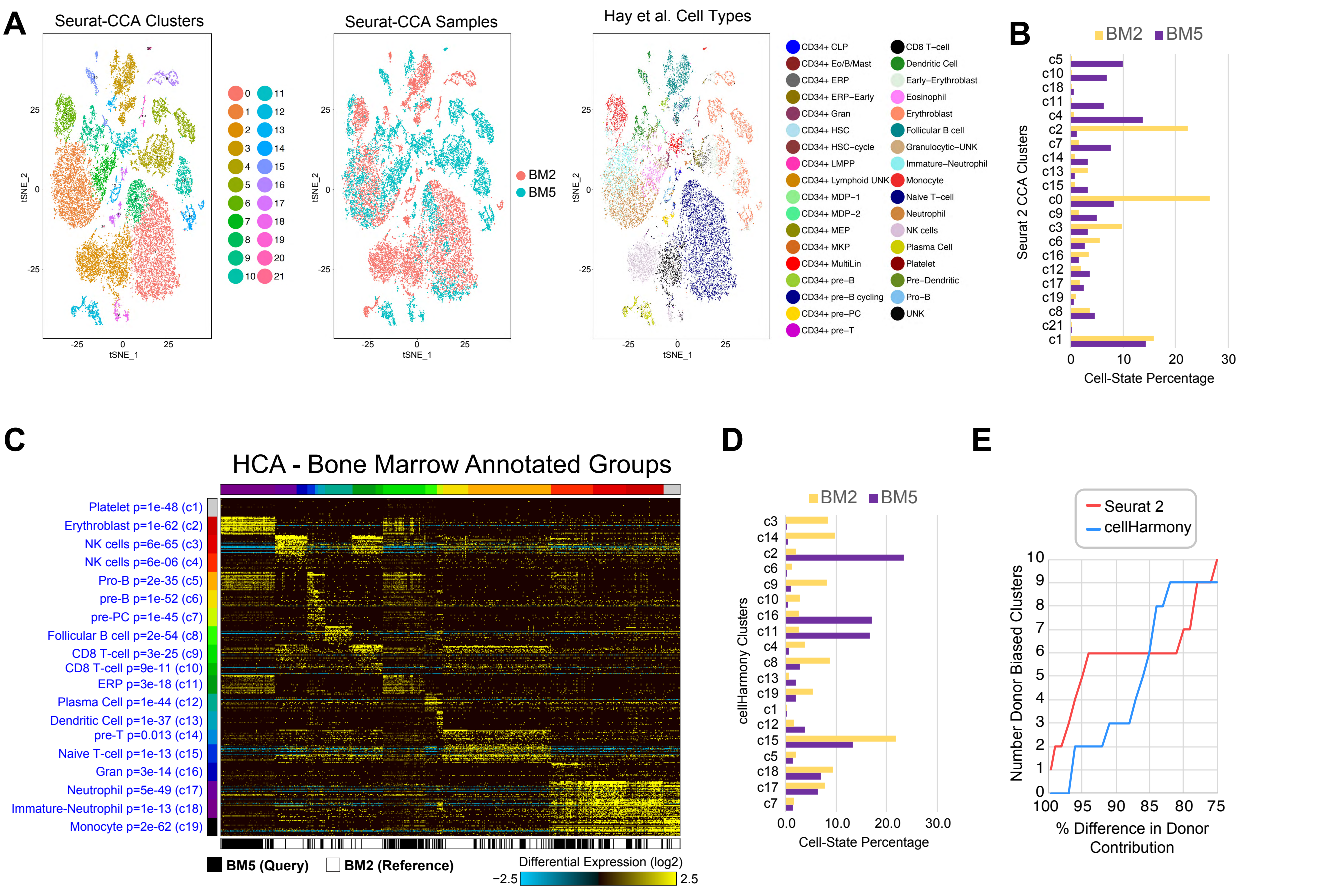

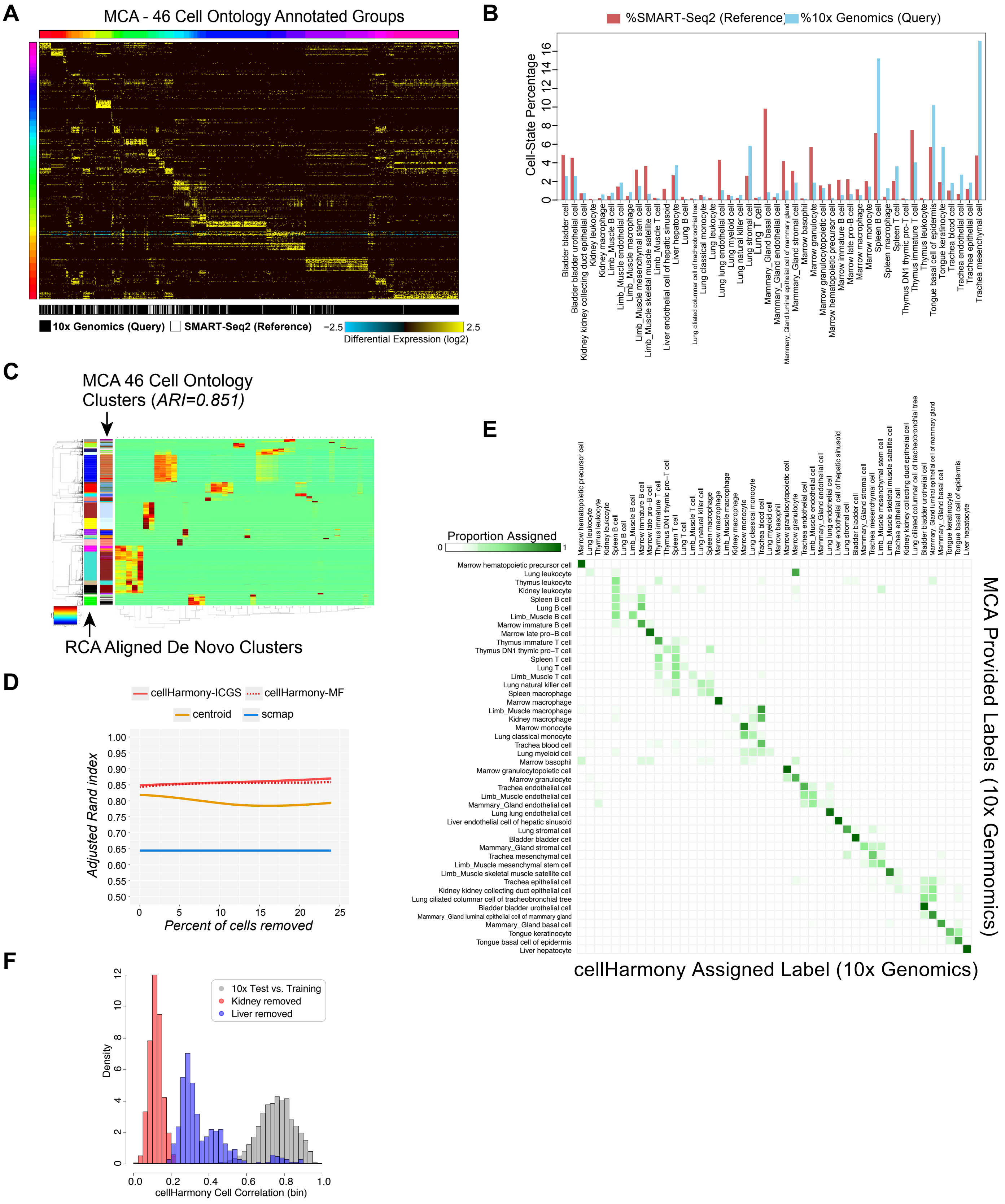

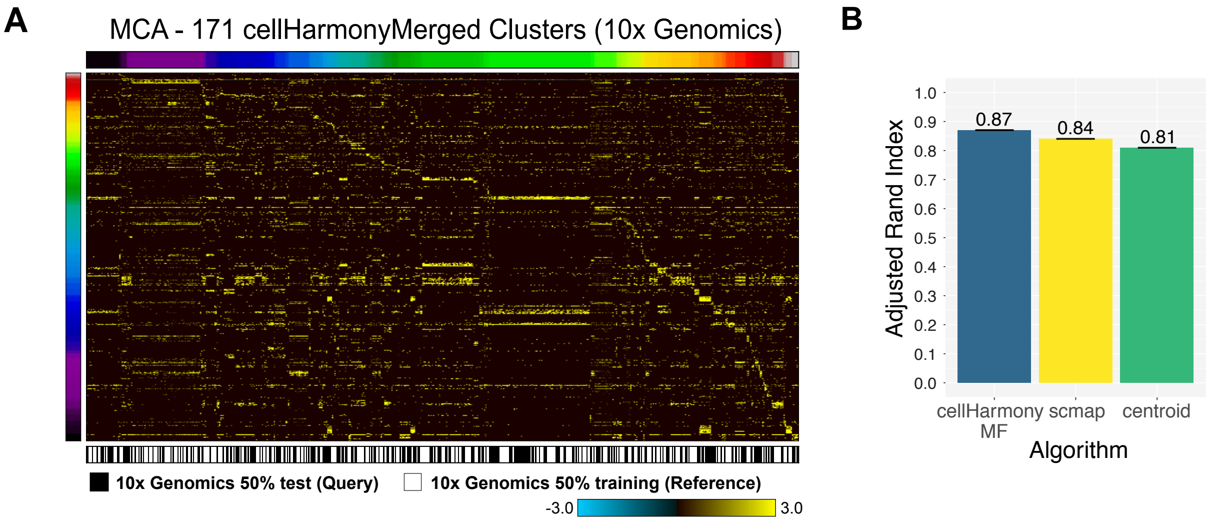

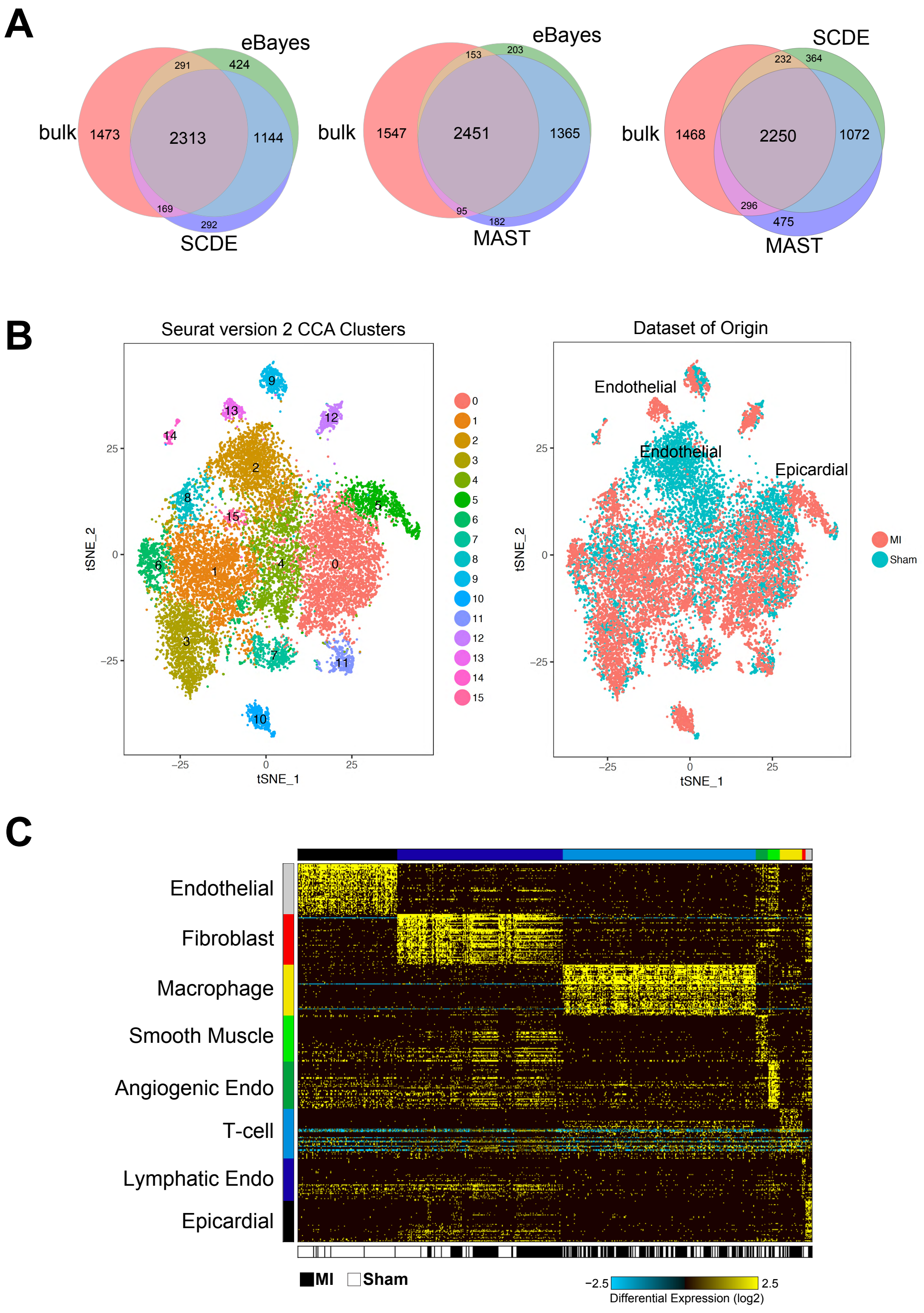

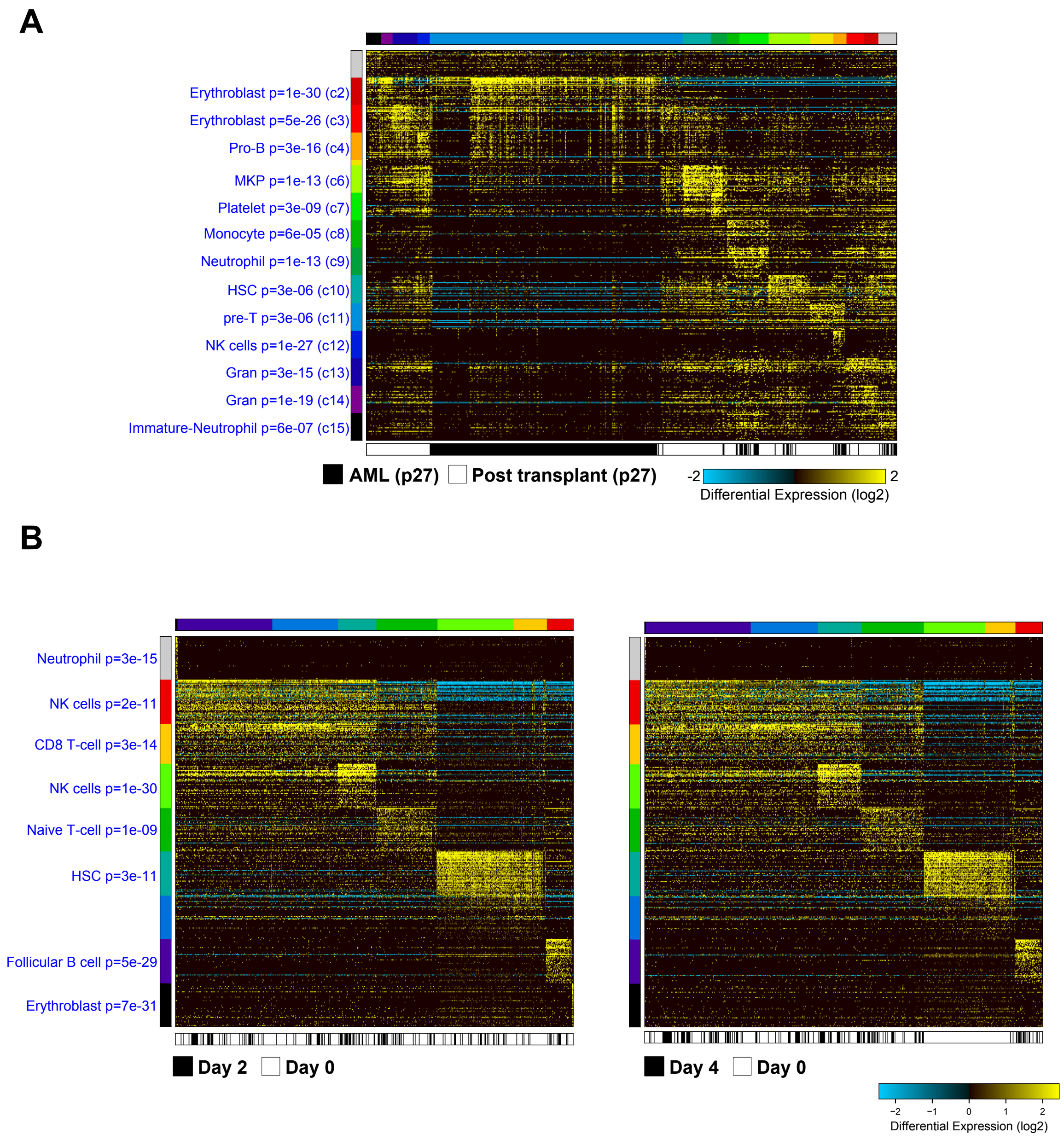
